# The structural basis of mRNA recognition and binding by eukaryotic pseudouridine synthase PUS1

**DOI:** 10.1101/2021.12.08.471817

**Authors:** Sebastian Grünberg, Lindsey A. Doyle, Eric J. Wolf, Nan Dai, Ivan R. Corrêa, Erbay Yigit, Barry L. Stoddard

## Abstract

The chemical modification of RNA bases represents a ubiquitous activity that spans all domains of life. Pseudouridylation is the most common RNA modification and is observed within tRNA, rRNA, ncRNA and mRNAs. Pseudouridine synthase or ‘PUS’ enzymes include those that rely on guide RNA molecules and others that function as ‘stand-alone’ enzymes. Among the latter, several have been shown to modify mRNA transcripts. Although recent studies have defined the structural requirements for RNA to act as a PUS target, the mechanisms by which PUS1 recognizes these target sequences in mRNA are not well understood. Here we describe the crystal structure of yeast PUS1 bound to an RNA target that we identified as being a hot spot for PUS1-interaction within a model mRNA at 2.4 Å resolution. The enzyme recognizes and binds both strands in a helical base-paired RNA duplex, and thus guides the RNA containing the target uridine to the active site for subsequent modification of the transcript. The study also allows us to show the divergence of related PUS1 enzymes and their corresponding RNA target specificities, and to speculate on the basis by which PUS1 binds and modifies mRNA or tRNA substrates.

## INTRODUCTION

The cellular transcriptome throughout all domains of life displays a highly complex regulatory network of more than 150 known post-transcriptional RNA modifications that modulate RNA biogenesis, function, specificity, and stability (1-3). Pseudouridine (Ψ), a C5-glycosidic isomer of uridine, was discovered in 1951 and soon after termed the fifth nucleoside (4-6). Almost 20 years later, the first pseudouridine synthase gene, *TruA*, was identified, and found to modify tRNA in bacteria (7,8). More recent advances in transcriptome-wide mapping of Ψ revealed widespread pseudouridylation of mRNAs in eukaryotes at levels comparable to m^6^A modifications (9-14). Despite the abundance of Ψ in mRNA, little is known about a) the purpose of that modification, b) whether the location of pseudouridine modifications in mRNA is random or if they are of functional importance, and c) whether they are installed by specific PUS enzymes.

Two unique classes of pseudouridine synthase enzymes, that are jointly responsible for pseudouridylation of RNA, differ in the way that they target their RNA substrates. Guide RNA-dependent PUS enzymes in eukaryotes and archaea (such as Cbf5 in yeast and dyskerin in humans) are part of a box H/ACA ribonucleoprotein complex and utilize small nucleolar RNAs (snoRNAs) as guides that recognize and base-pair with its substrates, which are mostly non-coding RNAs (ncRNA) (15). Alternatively, an RNA recognition mechanism that does not rely on guide RNA factors is employed by stand-alone PUS enzymes that independently recognize and modify their targets (reviewed in (16)). In contrast to the H/ACA snoRNP PUS enzymes, most stand-alone PUS enzymes are conserved throughout eukaryotes and bacteria. These PUS enzymes are further divided into six families, which differ in N- or C-terminal extensions flanking a conserved catalytic core domain. That domain is present in all H/ACA snoRNP and stand-alone PUS enzymes and contains a central catalytic motif corresponding to an antiparallel β-sheet that is flanked by α-helices (17,18). A strictly conserved catalytic aspartate residue in that motif is required for the rearrangement of uridine to its C-glycoside isomer Ψ in a mechanism that is not yet clear (19,20).

Eukaryotes generally contain multiple unique stand-alone PUS enzymes (PUS1-PUS10 and mitochondrial RPUSD), which differ in their substrate preference and localization in the cell. Each of them modifies specific sites in tRNAs, snRNAs, and rRNAs by targeting a sequence and/or structural element in their respective substrates (16,19). The crystal structures of many PUS enzymes have been solved, including structures of eukaryotic/archaeal box H/ACA snoRNPs with guide and substrate RNAs (21-28) and several bacterial stand-alone PUS enzymes bound to non-coding RNAs, most of them revealing specific interactions of PUS enzymes with their respective targets (29-32).

It remains somewhat unclear which PUS enzymes are responsible for the pseudouridylation of messenger RNA (mRNA). Various studies have indicated that more than half of the Ψ modifications in mRNA are catalyzed by the tRNA-specific PUS4 (a member of the divergent TruB family of PUS enzymes) and PUS7 (TruD family) through recognition of tRNA-like structures (which are their reported primary targets (10,12-14)). Like PUS7, PUS1 (TruA family) has been reported to modify multiple structurally diverse positions in tRNA, U2 and U6 snRNAs, suggesting less restricted selection of its RNA targets by these enzymes compared to PUS4 (33-36). More recent data have suggested that PUS1 is the predominant PUS to modify uridines in mRNA (33,37-39). Although PUS1 sites in mRNA show little sequence similarity, high-throughput pseudouridylation studies have implicated a structure-dependent mRNA target recognition mechanism and suggested that modulation of the RNA structure may play a role in the regulation of mRNA pseudouridylation (14,40). However, it is still unclear how a tRNA-specific pseudouridine synthase recognizes and modifies mRNA substrates, mostly due to the lack of available structural information of PUS1 enzymes bound to mRNA targets.

Only a few structures of eukaryotic stand-alone PUS enzymes have been solved: the catalytic domains of human PUS1 and PUS10 (41-43), as well as structures of human and yeast PUS7 (44,45). The only available structural information of a stand-alone PUS enzyme bound to its target RNA are the structures of *E. coli* TruA (the closest bacterial homologue of PUS1) and TruB, each bound to tRNA (30,46). Despite the structural similarities between TruA and PUS1, modeling and docking studies of tRNA and the core domain of human PUS1 suggests a significantly different orientation of the tRNA when bound to PUS1 than in the *E. coli* tRNA-TruA complex (43).

Because visualization of how PUS1 binds to mRNA might provide new insight into the basis of its action on such substrates, we generated a pair of crystal structures of wildtype *S. cerevisiae* PUS1 and a catalytically inactive PUS1 mutant in complex with short mRNA fragments. The structures, from two unrelated crystal forms, represent the first structures of a eukaryotic stand-alone PUS enzyme bound to a target RNA. The structures illustrate how PUS1 recognizes and binds a helical RNA duplex and enable the identification of RNA-contacting amino acid residues that make extensive contacts with each RNA strand to orient and guide the RNA into the active site of the enzyme. Additional examination and comparison with previously described structural and biochemical studies indicate that a) while PUS1 and TruA use the same protein surface to interact with their respective target-RNAs, the position and identity of the corresponding contact residues and the position of the bound RNAs differ significantly from one another, and b) PUS1 likely binds and acts on at least one tRNA target in a manner that is closely related to how it engages with its docking site in mRNA.

## MATERIALS AND METHODS

### Cloning, Protein Expression and Purification

N-terminally HIS-tagged *S. cerevisae* PUS1 (PUS1; NEB # M0526S) was used for most biochemical assays. A catalytically inactive PUS1 was generated by single point mutation of active site residue aspartate 134 into an alanine (PUS1_D134A_). For biochemical characterization, the mutant PUS1 was expressed for two hours at 37°C in T7 Express lysY competent *E. coli* (NEB # C3010). Cells were harvested, washed with cold 1x PBS, resuspended in HisTrap binding buffer (HTBB; 20 mM Na_2_HPO_4_, 0.5 M NaCl, 20 mM Imidazole, 1 mM DTT, 20% glycerol) + 1x protease inhibitors (1 mM PMSF, 0.5 nM Leupeptin, 2.75 mM Benzamidine, 2 nM Pepstatin) and stored at -20°C. Cells were thawed, sonicated, and after centrifugation, the supernatant was saved and applied to a 5 ml HisTrap HP column (Cytiva) calibrated with HTBB. PUS1_D134A_ was eluted with HisTrap elution buffer (20 mM Na_2_HPO_4_, 0.5 M NaCl, 0.5 M Imidazole, 1 mM DTT, 20% glycerol). Fractions containing PUS1 were pooled and dialyzed into HiTrap Heparin binding buffer (10 mM Na_2_HPO_4_, 100 mM NaCl, 20% glycerol, 1 mM DTT).

The dialyzed protein was applied to a 5 ml HiTrap Heparin column (Cytiva) and eluted with HiTrap Heparin elution buffer (10 mM Na_2_HPO_4_, 2 M NaCl, 20% glycerol, 1 mM DTT). The fractions containing PUS1 were pooled and dialyzed into HiTrap SP binding buffer (50 mM HEPES, pH 7.0 at 4°C, 50 mM NaCl, 1 mM DTT, 0.1 mM EDTA). The dialyzed PUS1 was applied to a 5 ml HiTrap SP HP cation exchange column and eluted with HiTrap SP elution buffer (50 mM HEPES, pH 7.0 at 4°C, 1 M NaCl, 1 mM DTT, 0.1 mM EDTA). Finally, fractions containing PUS1 were pooled and dialyzed into storage buffer (10 mM Tris/Cl pH 7.4 at 4°C, 200 mM NaCl, 1 mM DTT, 0.1 mM EDTA, 50% Glycerol, 200 µg/ml BSA) and stored at - 20°C.

To generate an untagged version of the PUS1 enzyme for crystallography trials, PUS1_D134A_ lacking the N-terminal HIS-tag was subcloned into expression vector pET21d (EMD Biosciences) using a Gibson Assembly cloning kit and protocol (NEB # E5510S) and sequence verified. A clone encoding tagless wild type PUS1 was then generated from PUS1_D134A_ by converting alanine 134 back to wild type aspartate 134 using a QuickChange II XL kit and protocol (Agilent).

Sequence-verified constructs were transformed into BL21-CodonPlus(DE3)-RIL competent cells (Agilent) and expressed using a previously described autoinduction protocol (47). Bacterial pellets from a liter of culture were resuspended in 30 mL of Buffer A (50 mM NaCl, 10 mM Tris pH 7.5, 0.1 mM EDTA, 5% glycerol, 1 mM DTT), and 0.2 mM PMSF and 900 U of Benzonase added. Cells were lysed on ice with a Misonix S-4000 sonicator operating at 70% power; the cell suspension was subjected to 80 seconds of total sonication time over the course of four cycles (each applying 20 second sonication bursts followed by 60 second cooling periods). The cell lysate was clarified by centrifugation in an SS-34 rotor (Sorvall) at 30597 x g for 20 minutes at 4°C, followed by hand filtration through a 5 Ιm filter. The lysate was loaded onto a 5 mL HiTrap Heparin HP column (Cytiva) and the column was washed with 25 mL of Buffer A. A gradient from 100% Buffer A to 100% Buffer B (Buffer A augmented with 1.0 M NaCl) was run with a total elution volume of 100 mL and 5 mL fractions were collected. Peak fractions were combined, concentrated, filtered, and loaded onto a HiLoad 16/60 Superdex 200 gel filtration column (Cytiva) equilibrated in Buffer SEC (Buffer A augmented with 200 mM NaCl). Peak fractions were combined and concentrated. The final purification of the protein, including the size exclusion chromatography elution profile of the purified protein, are shown in **Supplemental Figure S1.**

### *In vitro* transcription of Fluc mRNA fragments

A DNA duplex containing the T7 promoter followed by Fluc positions 787-845 was ordered from Integrated DNA Technologies (5’ - gcgaaattaatacgactcactatagggATTCCGGATACTGCGATTTTAAGTGTTGTTCCATTCCATCACGGTTTTGGAATGTTTAC – 3’). 1 µg of the duplex was *in vitro* transcribed using the HiScribe™ T7 Quick High Yield RNA Synthesis Kit (NEB # E2050S) following the standard IVT protocol and purified using Monarch® RNA Cleanup columns (NEB # T2030).

### PUS1 activity assays

210 pmol (2.1 µM) PUS1 was incubated with 84 pmol (0.84 µM) synthetic RNA oligos (Integrated DNA Technologies, Supplementary Table S1) or a short in vitro transcript (IVT) in 1x NEB buffer 1.1 for 90 min at 30°C in a 100 µL reaction. Reactions with full length Fluc mRNA contained 2.1 µM PUS1 and 8.3 nM Fluc mRNA). Reactions were stopped by adding 0.8 units of Proteinase K (NEB # P8107S) followed by incubation for 10 min at 37°C. The modified RNA was subsequently column-purified (NEB # T2030), 2 pmol were digested to single nucleosides using the Nucleoside Digestion Mix (NEB # M0649), and the ratio of Ψ s versus uridines was determined via tandem quadrupole mass spectrometry (Supplementary Figure S2a).

### Crystallization and Data Collection

RNA targets were complexed with purified protein, with the RNA present in a 1.4-molar excess over the protein. Complexes were screened for crystal grown in 96 well plate formats, with 200 nanoliter drop volumes equilibrated against 100 microliter reservoirs, against multiple commercial crystallization screens, while using a mosquito robot (TTP LabTech). Drops that generated visible crystals were then used to set up subsequent screening and expansion trays by hand, with 2 microliter drops equilibrating against 1000 microliter reservoir volumes.

A mixture of catalytically inactive PUS1 (PUS1_D134A_) at 11.6 mg/mL and a slight molar excess of RNA (‘R263’; 5’-AAA UCG GGA UUC CGG AUA-3’) crystallized at 4° C after equilibration against a reservoir containing 0.05 M Ammonium sulfate, 0.05 M Bis-Tris pH 6.0, and 26% Pentaerythritol ethoxylate. In contrast, a mixture of wildtype PUS1 (PUS1_wt_) at 11 mg/mL in the presence of a similar RNA construct harboring a 5-fluorouracil moiety at a position corresponding to potential enzymatic modification (‘R340’; 5’-[5FU]AA UCG GGA UUC CGG AUA-3’) crystallized at 25°C after equilibration against a reservoir containing 0.2 M Potassium sodium tartrate and 18% PEG 3350 at 25°C.

Crystals were transferred to a cryocooling solution containing either 0.04 M Ammonium sulfate, 0.04 M Bis-Tris pH 6.1, 23% Pentaerythritol ethoxylate, and 22% ethylene glycol or 0.2 M Potassium sodium tartrate, 20% PEG 3350, and 20% ethylene glycol for PUS1_D134A_ or PUS1_wt_, respectively. Crystal were then flash frozen in liquid nitrogen and data was collected on ALS BCSB beam line 5.0.1. Data for PUS1_D134A_ was processed using program HKL2000(48). Data for PUS1_wt_ was automatically processed using program XDS(49).

### Phasing and Refinement

Structures were phased via the molecular replacement method, using the catalytic domain of human PUS1 (PDB ID 4J37) (43) as a search model. Molecular replacement searches were conducted using program PHASER (50) within the Phenix crystallographic computational suite (51). The structure was rebuilt and refined using programs COOT (52) and PHENIX.REFINE (53).

## RESULTS

### Identification of a *PUS1 mRNA substrate*

Since we were interested in the ability of PUS1 to modify uridines in full length mRNA, we tested the activity of purified PUS1 on *in vitro* transcribed Firefly luciferase (Fluc; 1766 nt, **Supplemental Table S1**) mRNA, a standard model mRNA in our lab. Using LC-MS/MS to detect and quantify the number of uridines that were pseudouridylated by PUS1, we were surprised to find that approximately 14 % of all uridines in Fluc mRNA were pseudouridylated after 5 hours of incubation with PUS1 (**Supplemental Figure S2b**). These values suggested that approximately 64 of the total 458 uridines in Fluc had been modified by PUS1. These data did not provide information about whether the same uridines at specific positions in Fluc were modified, or if the pseudouridylation occurred in a non-targeted fashion and was distributed randomly across the mRNA. However, this number was significantly higher than the less-than-one Ψ per transcript that have been reported for cellular mRNAs previously (14).

Using a sliding window approach, dividing the Fluc mRNA first into 300 nt and then by 60 nt fragments, we identified a sequence spanning positions 787 to 845 to be highly pseudouridylated by PUS1 (**Supplemental Figure S2c**). To facilitate subsequent crystallographic studies, we further tested multiple short synthetic RNA oligos covering Fluc positions 760 – 845 to find the shortest possible mRNA substrate for PUS1 (**Supplemental Figure S2d**). We found that most oligos comprising the Fluc sequences between positions 760 and 796 displayed robust pseudouridylation, while oligos with sequences downstream of position 787 were only weakly or not modified at all (**Supplemental Figure S2e and S2f**). Concurrent to our work, the Gilbert lab combined computational prediction and mutational analysis to reveal the determinants for an RNA target to be recognized by PUS1. Their study suggested that the ideal PUS1 RNA substrate contains a uridine as part of a weak H-R-U motif at positions -2, -1, 1 at the base of a 13 nt/15 nt stem-loop structure (40). Interestingly, when we predicted the secondary structure of each of the RNA oligo substrates, we found that oligos that were modified by PUS1 all formed a suboptimal, but similar, PUS1 target structure (**Supplemental Figure S2g**). The pseudouridylated oligos R167, R168, and R169 all contained at least one uridine at the base of a stem-loop structure. However, R167 and R168 did not contain a H-R-U motif, and the stem length – a critical component in PUS1 target preference – in all oligos was shorter than optimal.

Intrigued by these results, we hypothesized that we could engineer an optimized RNA substrate following the parameters described by the Gilbert lab, but based on the Fluc mRNA sequence that would (1) be efficiently modified by PUS1, and (2) form stable complexes with PUS1 for crystallography (**Supplementary Figure S3a**). When we incubated the engineered Fluc stem-loop substrate (R397) with PUS1 *in vitro*, we identified pseudouridylation in an RNA fragment encompassing the 5’-end of the substrate via UHPLC-MS/MS (**Supplemental Figure S3d (panels 1, 2)**). To confirm the position of the pseudouridine site, we incubated PUS1 with a similar substrate (R444) featuring reversed positions of the U and A at the base of the stem (**Supplemental Figure S3b**). This substrate did not get pseudouridylated by PUS1 (**Supplemental Figure S3d, panel 3**), confirming that - as predicted - PUS1 only targeted the uridine in the H-R-U motif at the base of the stem. In addition, we designed an artificial RNA substrate (R398) that fused the R397 Fluc-mRNA stem loop with the known PUS1 target PFY1-U290 mutant stem 1 loop that had previously been shown to be pseudouridylated by PUS1 *in vitro* (40). As expected, we detected pseudouridylation in the fragments containing the H-R-U motif at the base of either stem (**Supplemental Figure S3c, d (panels 3 and 4)**).

### An active site mutant of PUS1 is catalytically inactive

We generated a catalytically inactive PUS1 variant, with the intention of using such a construct for structural studies without the complication of formation of a heterogeneous mixture of substrate, reactions intermediates or products during the crystallization process. To do so, we exploited the observation that all PUS enzymes – stand-alone or as part of the H/ACA snoRNP complexes – contain a conserved aspartate residue (D146 in human PUS1) that is strictly required for activity (43,54). Substituting the aspartate residue 134 with alanine (D134A) in yeast PUS1 generated a catalytically inactive PUS1 variant (PUS1_D134A_) that was unable to pseudouridylate RNA (**Supplemental Figure S4**).

### PUS1 binds to an RNA duplex

Even though both optimized RNA substrates R397 and R398 were recognized and pseudouridylated by PUS1 at the predicted positions, attempts to co-crystallize PUS1 with either substrate were unsuccessful, most likely due to steric interference of the long RNA extensions with crystal lattice formation. We thus attempted crystallization with shorter RNA oligo substrates that resembled suboptimal PUS1 target structures that we had identified as being efficiently modified (R167, R168, R169). Unfortunately, neither of these oligos formed crystals with PUS1_wt_ or PUS1_D134A_ either. We then focused on RNA oligo R263 (sequence provided in **Supplemental Figure S2f**) as potential substrate for co-crystallization with PUS1. While this short 18 nt oligo only displayed very low levels of pseudouridylation by PUS1 *in vitro* (**Supplemental Figure S2e**), it was predicted to form a short stem-loop structure with a uridine at the 5’ base of the stem, suggesting that it may still be recognized by PUS1 (**Supplemental Figure S2g**). When tested for PUS1-binding, R263 formed stable complexes with PUS1_wt_ and PUS1_D134A_ in electrophoretic mobility shift assays (EMSA) (**Supplemental Figure S5**). We therefore decided to attempt crystallization of both PUS1 enzymes in the presence of R263.

### PUS1 interacts with both strands of a helical RNA duplex

PUS1-RNA co-crystals that diffracted to 2.4 Å resolution, containing the R263 RNA construct and the PUS1_D134A_ inactive enzyme, were successfully grown and found to belong to space group C2 (**Table 1**). Modeling and refinement of the enzyme-RNA complex yielded values for the crystallographic R-factors (R_work_ and R_free_) of 0.222 and 0.265 respectively, with tight protein geometry (rmsd bonds and angles 0.003 Å and 0.58°; 95.73% of residues in favored Ramachandran regions). The average B-factor values for the protein and the bound RNA were comparable (65.48 and 62.74 Å^2^, respectively). The electron density maps from molecular replacement and subsequent rounds of refinement displayed well-ordered density for most of the protein chain (the first 70 residues, last 49 residues, and four subsequent surface loops ranging in length from 2 to 19 residues were disordered) and the first 12 bases of the RNA substrate (the final 6 base pairs were also disordered).

**Table 1.** Crystallographic Data and Refinement Statistics.

|  | scPUS1_D134A + sgR263 | scPUS1 + sgR340 |
| --- | --- | --- |
| *PDB ID* | *7R9G* | *7R9F* |
| **Data Collection** |  |  |
| Space group | C2 | P 6<sub>1</sub> 2 2 |
| Unit cell |  |  |
| a, b, c | 104.2, 98.3, 72.2 | 134.5, 134.5, 158.1 |
| alpha, beta, gamma | 90, 102.8, 90 | 90, 90, 120 |
| Wavelength (Å) | 0.9774 | 0.9774 |
| Resolution range (Å) | 50-2.4 (2.44-2.4) | 50-2.89 (2.993-2.89) |
| Unique reflections | 27637 | 19497 |
| R-merge | 0.072 (0.632) | 0.026 (0.730) |
| R-meas | 0.078 (0.698) | 0.036 (1.033) |
| R-pim |  | 0.026 (0.730) |
| CC1/2 | (0.813) | (0.502) |
| Mean I/sigma(I) | 22.2 (1.6) | 15.62 (1.09) |
| Chi^2 | 0.74 |  |
| Multiplicity | 6.5 (5.3) | 2.0 (2.0) |
| Completeness (%) | 99.6 (95.1) | 99.9 (99.84) |
| Wilson B-factor | 53.3 | 93.4 |
| **Refinement** |  |  |
| R-work | 0.2217 | 0.2537 |
| R-free | 0.2647 | 0.2964 |
| Number of non-hydrogen atoms | 3234 | 2962 |
| macromolecules | 3178 | 2938 |
| ligands | 6 | 24 |
| solvent | 50 | 0 |
| Protein residues | 385 | 379 |
| RMS(bonds) | 0.003 | 0.004 |
| RMS(angles) | 0.58 | 0.75 |
| Ramachandran favored (%) | 95.73 | 89.95 |
| Ramachandran allowed (%) | 3.47 | 7.34 |
| Ramachandran outliers (%) | 0.8 | 2.72 |
| Clashscore | 6.51 | 9.17 |
| Average B-factor | 65.36 | 91.61 |
| macromolecules | 65.48 | 91.63 |
| ligands | 62.74 | 88.32 |
| solvent | 58.11 | N/A |
| Maximum-Likelihood Coord. Error (Å) | 0.29 | 0.49 |

The contents of the crystallographic asymmetric unit correspond to a single protein subunit and a single strand of the bound RNA ligand (**Figure 1a**). That complex was observed to be part of a higher-order assembly, comprised of an RNA duplex and two bound copies of PUS1, that is generated via application of a crystallographic dyad symmetry axis (**Figure 1b**). Rather than binding to PUS1 as a monomer in the predicted short stem-loop formation, two R263 oligos formed a helical duplex. The base pairs within the RNA duplex (involving positions 2 through 10 in each strand) display both Watson-Crick and non-Watson-Crick interactions with their counterparts, including two central G:G reverse Hoogstein basepairs. The 5’- and 3’-most modeled base on each strand are not engaged in base-paired interactions but are also not flipped out of the strand’s duplex conformation (**Figure 1c**). Immediately proximal to the 5’ end of each RNA strand, at a distance of approximately 5.5 angstroms, a well-occupied and tightly coordinated sulfate ion is observed. This corresponds to a location and distance appropriate to represent an additional backbone phosphate group if the RNA were extended by an additional base at its 5’ end (**Figure 1d**). A sulfate ion was also observed and modeled at the same position in the previously described structure of the human PUS1 apo enzyme (PDB 4J37) (43).

**Figure 1.**
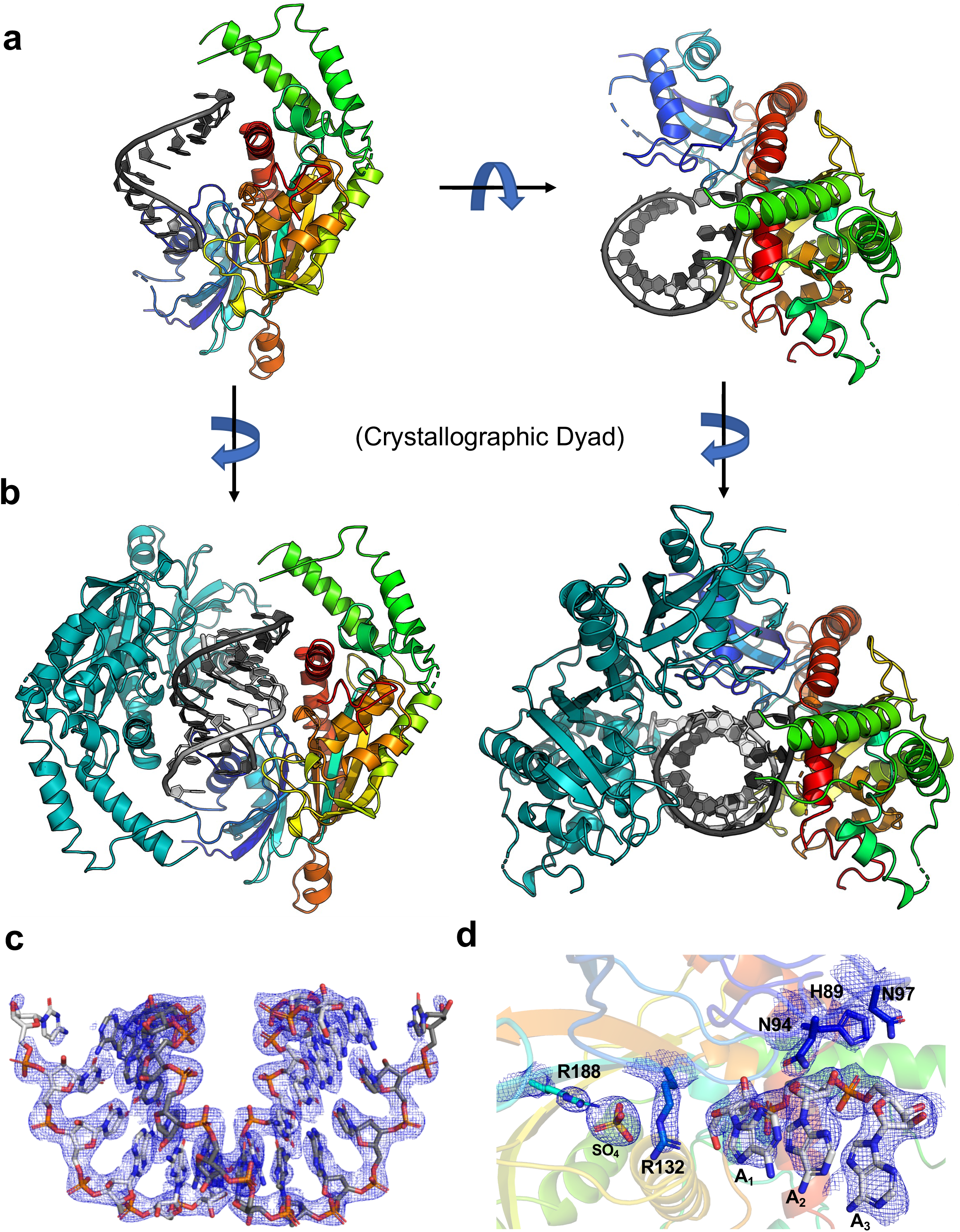
Model and representative electron-density for RNA-bound PUS1. ***Panel a:*** The contents of the asymmetric unit, for both structures that were solved, corresponds to a single protein subunit (colored as a spectrum, from the blue N-terminal end of the refined model to the red C-terminal end) bound to a single RNA oligonucleotide (grey bases). The model of the catalytically inactive D134A enzyme is shown in two orientations related by a 90° rotation around the x-axis. ***Panel b:*** In both structures, the application of a crystallographic 2-fold dyad rotation axis generates a dimeric complex in which two subunits are independently bound to an RNA duplex. The second protein subunit and second RNA strand are colored in dark teal and lighter grey, respectively. ***Panel c:*** Representative simulated annealing composite omit 2Fo-Fc electron density contoured across the RNA duplex and ***Panel d:*** at the region of protein-RNA contacts observed at the 5’ end of one RNA strand. The structural features illustrated in this figure are replicated for the wild type enzyme bound to a closely related RNA complex, which was solved in an unrelated crystallographic space group and lattice (**Supplementary Figure S9**). The position of the bound sulfate ion mirrors a similarly placed sulfate in the previously described structure of unbound human PUS1.

In contrast to the extensive base-paired contacts between the two crystallographically related RNA strands, the corresponding pair of bound protein molecules do not display significant contacts with one another; the few contacts between the two bound protein subunits are limited to two surface-exposed loops (residues 86 to 89 and 95 to 98) in the N-terminal region of the enzyme. Therefore, we believe that the two copies of PUS1 that are associated with opposite sides and ends of the symmetric RNA duplex are bound as individual monomers, independently of one another. That conclusion agrees with the solution behavior of the purified enzyme, which eluted from a size exclusion column as a monomeric species at high micromolar concentrations of protein (**Supplemental Figure S1**).

Within the complex between a single bound protein subunit and the RNA duplex, at least thirteen amino acid side chains contact numerous atoms on each of the two RNA strands (**Figure 2a and b**). The residues involved in substrate recognition and binding are largely comprised of two separated clusters of residues within the enzyme’s sequence and structure. The first cluster of seven RNA-contacting amino acids is distributed between residues 89 and 188; they collectively contact one end of the RNA duplex and the adjacent sulfate ion, positioning the first base at the 5’-end of an RNA strand near the entrance to the active site. All but one of those residues are conserved across eukaryotic PUS1 homologues from yeast to humans (**Supplementary Figure S6**). A second cluster of four additional RNA-contacting amino acids is distributed between residues 362 and 394; they contact a series of bases and backbone atoms further downstream on one of the two RNA strands and are conserved across eukaryotic PUS1 enzymes (**Figure 2a and b**, **Supplemental Figure S6**). Finally, two additional residues (K277 and Y459) contact the RNA backbone near the opposite end of the bound RNA duplex. The yeast specific K277 is not conserved across eukaryotic PUS1 enzymes. While the tyrosine in position 459 is yeast specific, other eukaryotic PUS1 variants contain a conserved threonine in its respective position. All the protein-RNA contacts described above are duplicated via symmetry by the second RNA-bound protein subunit (**Supplemental Figure S7**); each independently bound protein monomer makes multiple contact to nucleotide bases and to backbone phosphate groups on both RNA strands.

**Figure 2.**
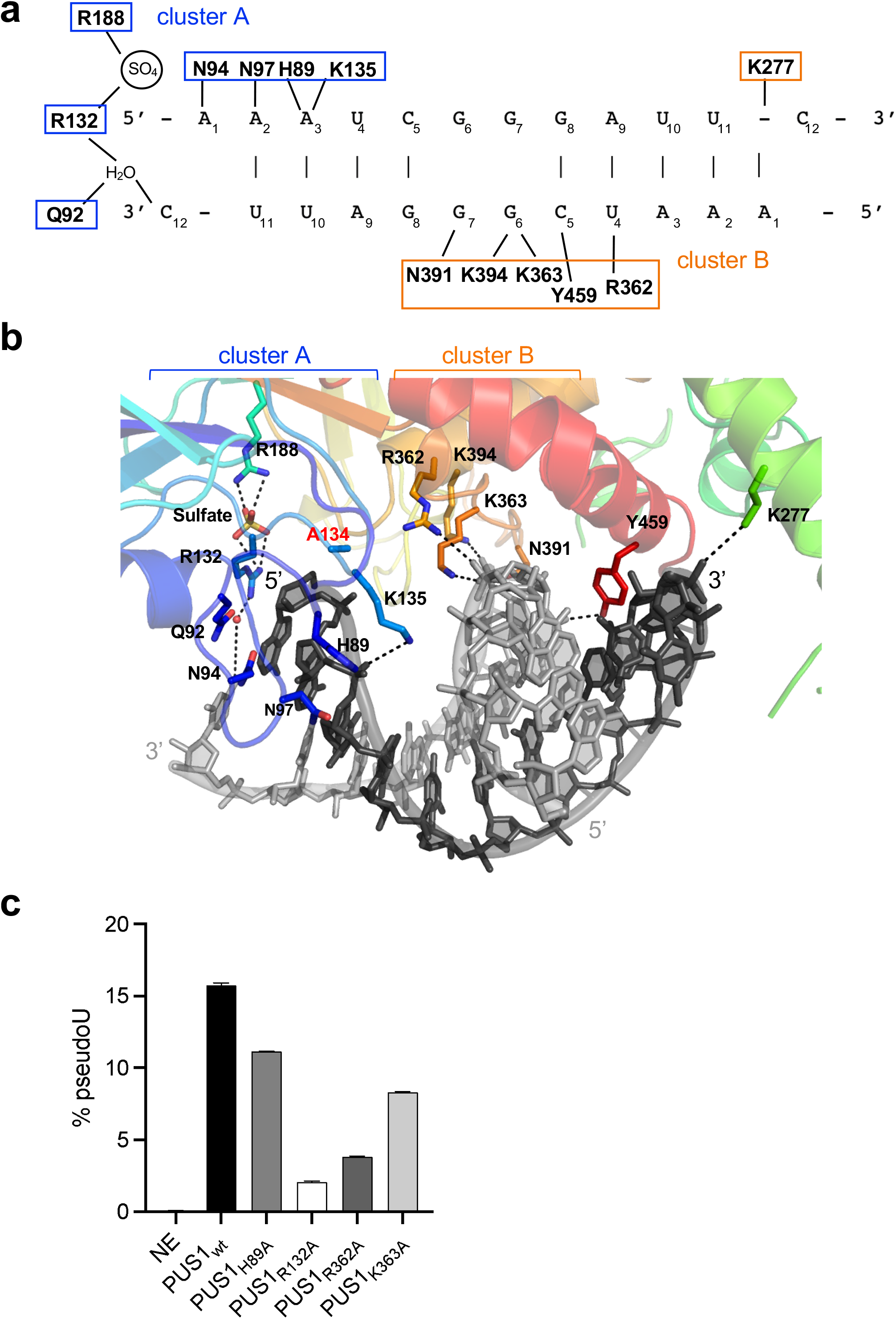
Protein-RNA contacts. ***Panel a:*** The first twelve nucleotides of each RNA strand are visible in the crystal structure, while the last six nucleotides are disordered and unobservable. Each protein subunit displays identical contacts, related by two-fold symmetry, to RNA bases and backbone atoms, and to a tightly coordinated sulfate ion immediately upstream of the upstream base in each bound RNA (contacts made by only one protein subunit are displayed for clarity). Residues that either contact the RNA near the active site (cluster a, blue boxes), or further downstream (cluster b, orange boxes) are indicated. ***Panel b:*** Distribution of RNA-contacting residues in the protein-subunit interface for one PUS1 subunit. Brackets indicate the residues that contact the RNA close to the active site (cluster A, blue) or further downstream (cluster B, orange). ***Panel c:*** Mutation of RNA-contacting amino acid residues in PUS1 negatively affects activity. The rate of pseudouridylation (% pseudoU) of PUS1 wildtype and the indicated PUS1 mutants with R169 substrate is shown. Data of two replicas is shown.

To further investigate if the identified protein-RNA contacts are important for PUS1 function, we generated alanine mutants of residues in either cluster A (H89, R132) or cluster B (R362, K363) that are in close proximity to the RNA strands. While replacing histidine 89 with alanine only had a weak effect on PUS1 activity and/or specificity when tested with the R169 RNA oligo (∼ 29% reduction of activity), mutating arginine 132 to alanine dramatically reduced pseudouridylation of the R169 RNA oligo by approximately 87%, suggesting an important functional role of this amino acid side chain in coordinating the RNA in the active site of the enzyme (**Figure 2c and Supplemental Figure S8a**). Despite not being in direct vicinity of the enzyme’s active center, residues in cluster B also proved important for PUS1 activity: Changing arginine 362 to alanine resulted in a dramatic reduction of pseudouridylation by approximately 75%, while mutation of the adjacent lysine 363 lead to a moderately reduced activity (∼ 47%; **Figure 3c and Supplemental Figure S8a**). To ensure that the differences in activity were not caused by varying PUS1 concentrations, we showed that each reaction indeed contained similar PUS1 concentrations (**Supplemental Figure S8b**). We then tested the wild type and mutant PUS1 variants with the engineered R397 substrate that contained only one Ψ-site. In agreement with the results above, we observed a dramatic reduction of pseudouridylation of the R397 RNA substrate with PUS1 mutants R132A and R362A, while PUS1H98A and PUS1K363A still modified their substrate (**Supplemental Figure S8c and b**).

**Figure 3.**
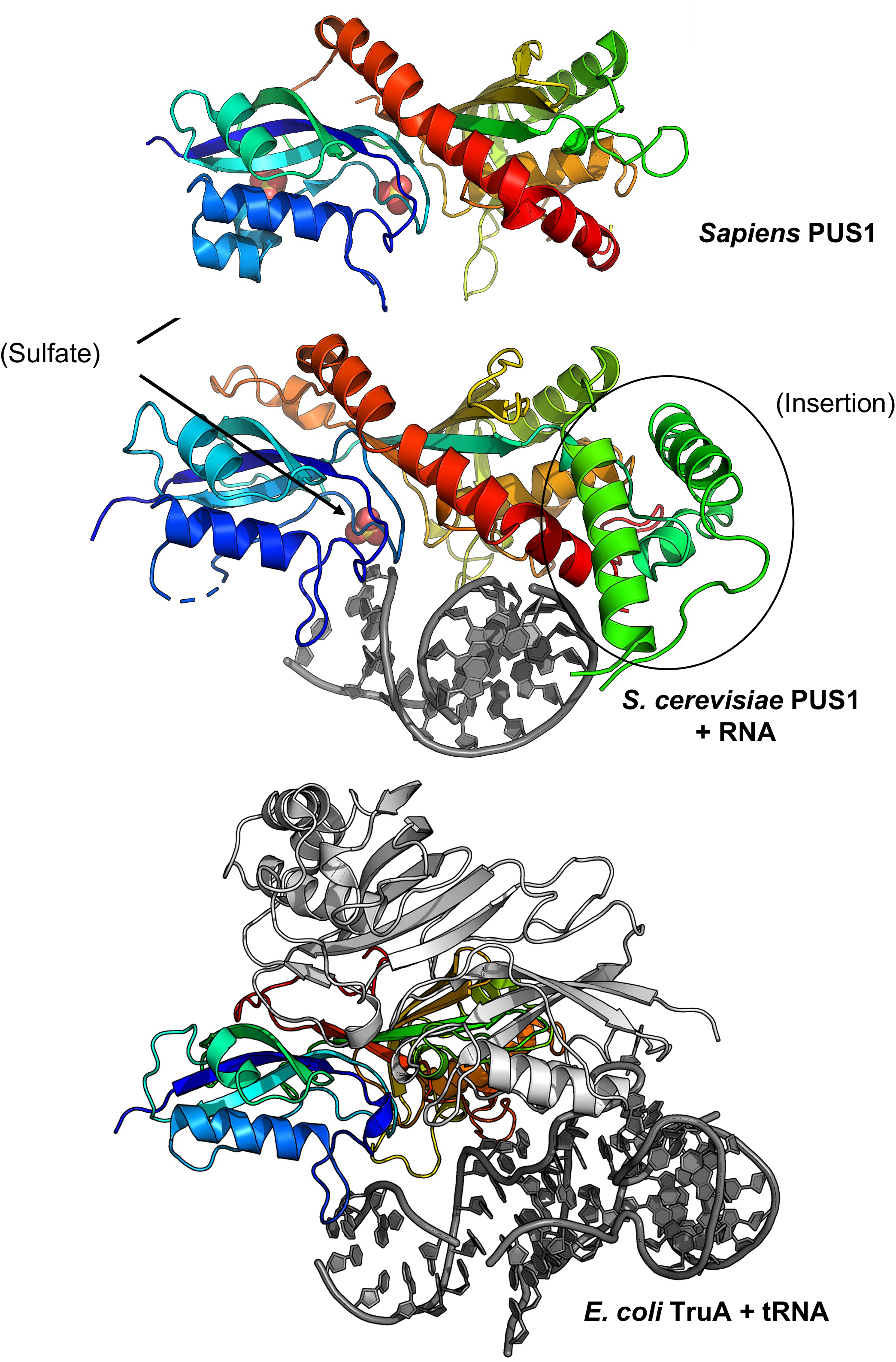
Comparison of yeast PUS1 to *E. coli* TruA and human PUS1. ***Panel a:*** Human PUS1 apo-enzyme (4J37). The structure includes two bound sulfate ions, one of which aligns with a single sulfate ion near the enzyme active site that was also observed in RNA-bound *S. cerevisiae* PUS1. ***Panel b***: *S. cerevisiae* PUS1_D134A_ bound to duplex RNA. A large insertion in the yeast enzyme, spanning residues S206 to approximately L279 (indicated by the oval), is unique as compared to its homologues in other eukaryotes (**Supplementary Figure S4**). It contains two RNA-contacting residues that are unique to the yeast enzyme. ***Panel c:*** *E. coli* TruA (2NR0) bound to tRNA. TruA utilizes an equivalent surface to bind its respective target but in a considerably different manner from PUS1.

To test whether mutations that significantly reduced PUS1 activity affected substrate binding, we performed EMSAs of wildtype and mutant PUS1 enzymes with the RNA substrate used to generate the crystal structures (R263) as described in Materials & Methods. Neither of the catalytically impaired cluster A and B mutants showed a reduction of substrate binding **(Supplemental Figure S8e)**. While these assays may not provide the resolution to accurately compare binding affinities, they clearly indicate that the detrimental effect of these mutations on enzyme activity cannot be explained by reduced binding. Together, these data imply that the formation and recognition of an RNA duplex near the active site is a mechanistically relevant feature of target selection.

In the structure of the catalytically inactive PUS1_D134A_ bound to the R263 duplex, the unpaired 5’ base (A_1_) is positioned within 4.5 angstroms of residue 134 (measured from C-alpha of 134 to C4’ of A1) and appears to be within potential distance to rotate into the enzyme active site (**Figure 2b**). To further examine the interactions of bound RNA with a catalytically competent version of the enzyme (in which the catalytic aspartate at position 134 was restored) we altered the RNA ligand by substituting a 5-fluoro-uracil (5-FU) base at position number 1, reasoning that it might be captured in a suicide complex by the enzyme after being flipped into the active site and subsequent nucleophilic attack by the carboxylate of D134. RNAs harboring such substitutions have previously been demonstrated to act as mechanism-based inhibitors of pseudouridine synthase enzymes and have been used to demonstrate the structural mechanism by which the target base gains access to the enzyme’s active site (55).

Crystals of wild type PUS1 in complex with the new R340 RNA construct (5’-[*5FU*]AA UCG GGA UUC CGG AUA-3’) belonged to a different space group (P6_1_22) and diffracted to 2.9 Å resolution. Although the space group and corresponding lattice packing arrangement of the protein-RNA enzyme complex was completely unrelated to the packing of the C2 space group of the PUS1_D134A_:RNA complex, the same RNA duplex, and the same position of two independently bound protein subunits was observed (**Supplementary Figure S9a**). The RNA duplex and second bound protein molecule is again generated by the application of a crystallographic dyad axis on the contents of the asymmetric unit (which again corresponds to a single protein subunit and a single RNA chain). Structural superposition of an RNA-bound monomer or dimer from the two crystal structures produced rmsd values of 0.63 Å and 1.47 Å, respectively. The reproducibility of all structural observations described above, in two different crystallization conditions and crystal forms, reinforces the conclusion that the formation and presence of an RNA duplex near the target uridine, and recognition of elements within each RNA strand by residues from each protein subunit, is a reproducible and mechanistically relevant feature of substrate recognition and activity.

Beyond providing an independent confirmation of the binding of two copies of the enzyme to an RNA duplex in a symmetric arrangement, the resulting electron density map, after several initial rounds of rebuilding and refinement, displayed a significant feature of positive difference density in the active site that might be indicative of low occupancy (ultimately estimated at less than 10% in refinement) by the 5’ 5FU nucleobase of the RNA (**Supplementary Figure S9b**). However, the combination of lower resolution and mixture of density features surrounding that base prevented us from unambiguously modeling the bound RNA in a state corresponding to a trapped catalytic complex. Interestingly, our attempts to modify the RNA substrate to trap either the target uridine or 5FU in the active site by adding different H-R-U motifs 5’ of the R263 sequence captured in our structures failed, despite designing the sequence to assure that the H-R part was single stranded.

The structure of the RNA-bound PUS1 enzyme was found to be similar to both the previously solved structure of human PUS1 in the absence of bound RNA (PDB 4J37) and the structure of bacterial tRNA pseudouridine synthase TruA (PDB 2NR0), with a rmsd across all comparable alpha carbons of approximately 2 Å in both pairwise superpositions (**Figure 3**). PUS1 and TruA employ the same protein surface, spanning the majority of their primary sequences, to contact their respective RNA targets; however, the position and identity of the corresponding contact residues and the conformation and orientation of their bound RNAs differ significantly from one another (**Supplementary Figure S10**). An alignment of the yeast PUS1 and *E. coli* TruA and their respective RNA substrates revealed that the orientation of the TruA-bound leucyl tRNA results in a clash of the tRNA with a yeast specific large insertion, spanning residues S206 to approximately L279 (**Figures 3, 4**).

**Figure 4.**
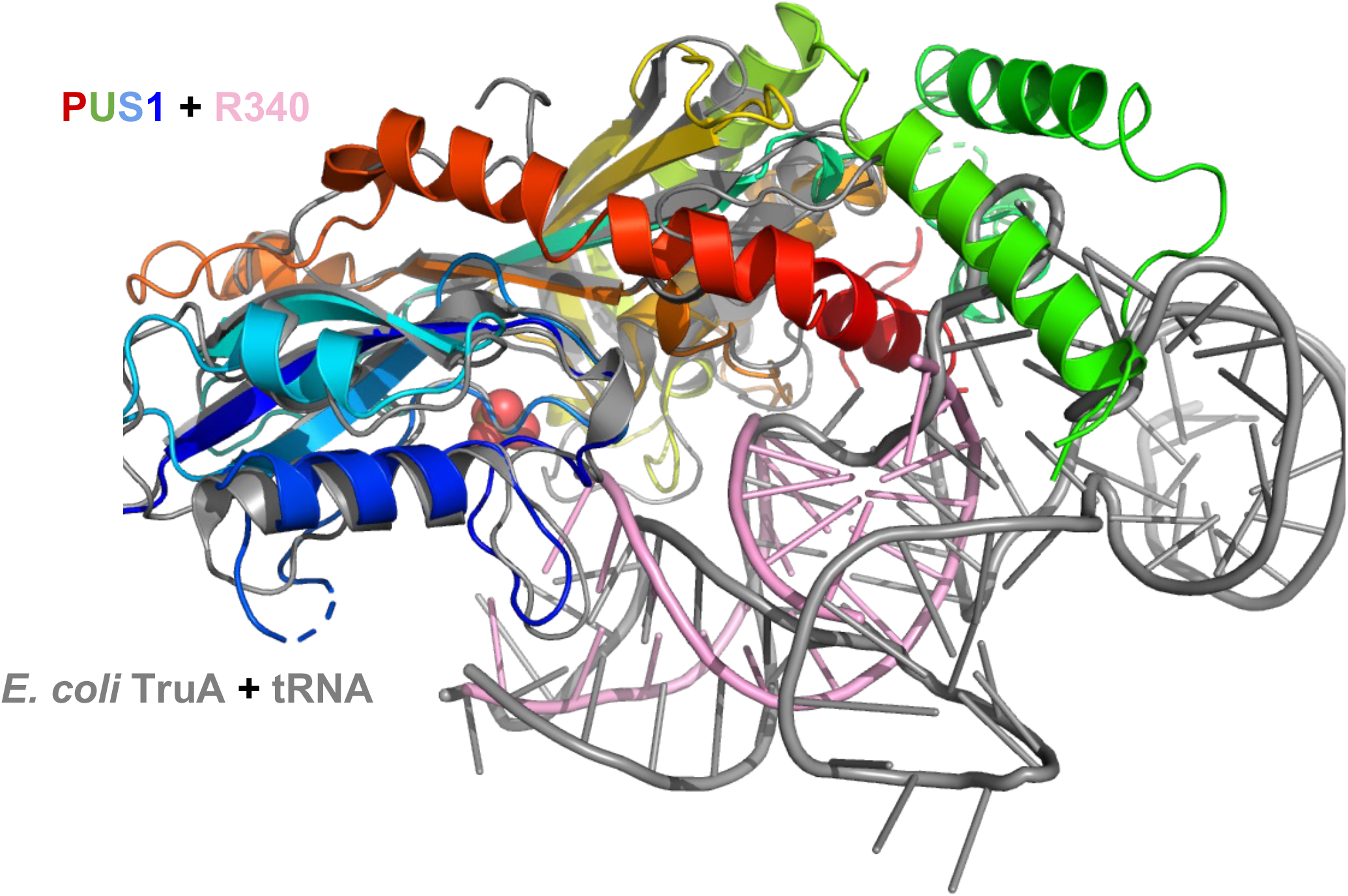
RNA-substrate binding by PUS1 and TruA. Superposition of RNA-bound PUS1 (PUS1: rainbow, RNA: pink) with tRNA-bound *E. coli* TruA (PDB: 2NR0; TruA & tRNA: gray) revealed a different orientation of the RNA substrates.

## DISCUSSION

Previous biochemical studies have provided solid evidence indicating how PUS enzymes might target and interact with their ncRNA substrates. However, it is still unknown how stand-alone PUS enzymes interact with mRNA. In addition, it is still unclear if PUS1-mediated mRNA pseudouridylation is site specific and functional, or non-targeted and mostly a side product of PUS1 non-specifically recognizing tRNA-like structures in mRNA. Our data present crystal structures of a guide RNA-independent eukaryotic PUS enzyme bound to a mRNA fragment and shed light on the molecular mechanism underlying mRNA recognition and binding by such enzymes.

While bacterial TruA has been reported to crystalize as a homodimer (with each subunit binding an individual tRNA), other PUS enzymes – including human PUS1 (41,43) – function as monomers. Therefore, the recruitment of two PUS1 to the RNA-duplex was initially surprising. However, our data and the crystal structures (solved in two different space groups) collectively indicate that yeast PUS1 does in fact interact with RNA substrates as a monomer, and that the observation of two bound enzyme subunits to the RNA duplex in a symmetric manner is a (fortuitous) biochemical and crystallographic artifact. First, the PUS1 enzyme is a monomer in solution as confirmed by size-exclusion chromatography (SEC) at high protein concentrations (**Supplemental Figure S1**). Second, despite binding to opposing ends and sides of the highly symmetric RNA-duplex used for crystallization, the PUS1 subunits do not display an extensive interface that would be expected for a functional protein dimer. However, we cannot exclude the possibility that binding of PUS1 to a single stranded RNA is required for or supports the formation of a stable RNA-duplex for some RNA substrates.

A question that immediately arises from our analyses is if the base-paired RNA duplex observed in the crystal structures represents a physiological and/or mechanistical state that influences PUS1 specificity and activity on mRNA. The Schwartz laboratory recently reported the importance of double-stranded stem RNA-structures for mRNA pseudouridylation by TruB1 and PUS7 through high-throughput Ψ mapping (10). Their data indicate that both enzymes modify structure motifs in mRNA that match important motifs in their preferred tRNA-substrates, suggesting a similar mechanism of site-recognition.

In contrast, PUS1 and its bacterial counterpart TruA are promiscuous in their target selection, modifying multiple RNAs with divergent sequences in the region of modification (56). Illumina-based mapping of Ψ installed by PUS1 in yeast and human cells revealed a weak H-R-U sequence motif, which the authors concluded was not sufficient to explain PUS1 specificity (14). However, a following study by the same laboratory reported that an RNA-duplex that forms a stem-loop structure in combination with the H-R-U motif is required for efficient pseudouridylation (40). They showed that features like stem-length and -stability regulate the pseudouridylation rate, which agrees with previous data showing that a loop flanked by stem structures is preferably targeted by human PUS1 (41). It also agrees with our results testing two stem-loop substrates. As predicted, only the uridines located at the base of the stem were pseudouridylated in both tested substrates. However, neither of the substrates formed crystals with PUS1. Despite adding a bulge in the stem region to increase conformational flexibility, the loop connecting both RNA strands may restrict the flexibility and the ability of the substrate to form its most stable orientation when forming a complex with PUS1 under the conditions used for crystallization. Interestingly, we observed PUS1 activity on short RNA oligos that were predicted to fold into structures with sub-optimal PUS1 target sites. 11% or 13% of 10 uridines in the R167 and R168 substrates, respectively, as well as 17% of the 12 uridines in R169 were pseudouridylated by PUS1 *in vitro*. Assuming that pseudouridylation occurred at specific sites rather than randomly throughout the oligo, this suggests that either one (R167/R168) or two specific uridines (R169) were modified by PUS1. Accordingly, PUS1 substrate selectivity – at least *in vitro* - may be less restricted than recently reported.

The crystal structure presented here provides first mechanical insight into why PUS1 requires a double stranded RNA stem for target recognition, binding, and activity. Our data show that each individual PUS1 subunit makes extensive contacts to both strands in the RNA duplex. Those RNA-contacting residues, which are well-conserved across PUS1 homologues, appear to enable binding of an RNA double helix and position the 5’ flanking region of the bound RNA strand close to the active site in PUS1. We find that alanine mutations of PUS1 residues arginine 132 and arginine 362 significantly reduce PUS1 activity. Interestingly, these two residues contact opposite strands in the RNA double helix, further supporting the hypothesis that an RNA double helix structure is important for binding of the RNA substrate and/or the positioning of the pseudouridylation site in the active site of the enzyme.

One explanation for yeast PUS1’s less restricted mode of structure recognition could be the location of its thumb loop (Tyr 83 – Thr 99). In the PUS1:mRNA structure, the loop is ordered and pointing away from the active site. This orientation results in a significantly wider opening of the active site cleft compared to TruA or human PUS1(43), which may allow for the binding of more diverse RNA substrates.

In agreement with our finding that PUS1 does not significantly modify the R263 substrate, none of the uridines within the crystallization oligonucleotide are located near the enzyme’s active site in the crystal structures but are instead involved in duplex formation via base pairing interactions between RNA strands. The structure suggests that an RNA-duplex with a minimum length of 10 to 12 nt is required for all RNA:PUS1 interactions to stably form and to position the RNA correctly in the active site. This agrees with a 11 bp median length of double stranded stem structures found in PUS1 mRNA targets (40). Our PUS1-dsRNA structures suggest that this minimum RNA-length is required for the formation of a sufficient stem structure that allows for all the PUS1-RNA interactions to form, which in turn positions the target uridine at the base of the stem in the catalytic center of the enzyme. Carlile and colleagues discuss the possibility that a cap at the entrance of the RNA binding channel in a yeast PUS1 structure model formed by a bundle of three α-helices may restrict the length of the RNA stem that can be accommodated in the enzyme (40,43). While our structures clearly indicate the presence of such a helical bundle, they do not indicate any clashes of the RNA with PUS1 components. Instead, we hypothesize that longer structures may not be efficiently bound, stabilized, and oriented in the active center by the PUS1-RNA interactions.

Comparison of the structures solved in this study with that of bacterial tRNA pseudouridine synthase TruA (30) (a representative of the structural family from which PUS1 is derived) bound to its tRNA substrate indicates how RNA recognition mechanisms can - and have - diverged dramatically. Both enzymes display contacts to their respective RNA targets involving approximately 15 residues distributed across their N- and C-terminal domains, and in doing so position their active sites near bases that are potential sites of modification (individual bases located 5’ of the bound RNA duplex for PUS1; bases 38, 39 and 40 of the tRNA anticodon stem loop for TruA). Of the residues involved in direct RNA contacts from each enzyme, several equivalent positions in each enzyme’s N-terminal region contact RNA atoms but involve quite different protein residues (<u>H</u>_89_GM<u>Q</u>Y<u>N</u>PP<u>N</u>_97_ in PUS1; <u>Y</u>_24_GW<u>Q</u>R<u>QNEV</u>_32_in TruA) that contact unique nucleotide identities and conformations in their corresponding RNA targets. In addition, the orientation of the TruA-bound leucyl tRNA causes a significant clash with two helices in a yeast specific insert (S206 - L279), providing further evidence for divergent evolution of these enzymes.

Conversely, a separate examination and comparison of the structures solved in this study against a tRNA known to be modified within its anticodon loop by PUS1 (33,43) indicates that the enzyme may be able to do so through a binding mode and interactions with that substrate’s anticodon stem loop that are similar to the binding and interactions observed in our crystal structure. Superposition of the tRNA substrate, via its anticodon stem loop, onto the RNA duplex from the crystal structure positions a base in the tRNA anticodon loop that is known to be modified by the enzyme immediately adjacent to the enzyme’s active site (**Figure 5**). The corresponding docked position of the tRNA further positions the previously identified RNA-contacting residues at similar distances and potential interactions with atoms along the tRNA backbone. The superposition does not indicate significant clashes between the C-terminal helical region of the PUS1 enzyme and the remainder of the tRNA molecule.

**Figure 5.**
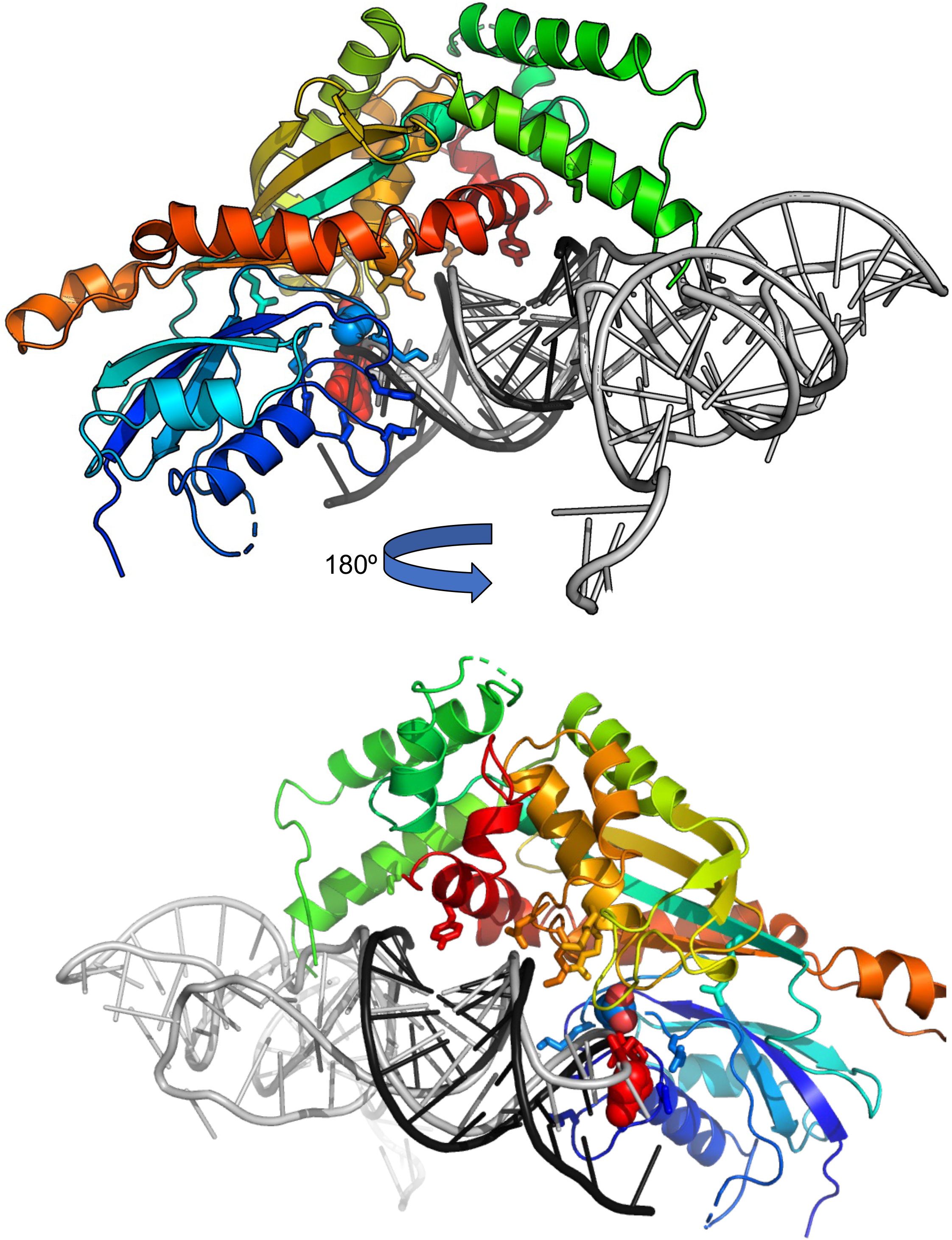
Superposition of the crystal structure of PUS1 bound to the RNA duplex described in this study with a human tRNA known to be modified by the same enzyme within its anticodon loop. The protein is colored; the RNA duplex from the crystal structure is dark grey; the tRNA substrate that is docked onto the crystal structure is light grey. The position of the active site D134 residue is indicated with light blue spheres; the position of the site of uracil modification is indicated with red spheres.

The structures presented here provide the first comprehensive mechanistic insight into the interaction between a eukaryotic stand-alone PUS enzyme and its mRNA substrates. However, some questions remain unanswered. First, is the underlying RNA sequence within a given RNA stem loop in an mRNA important for pseudouridylation activity? Or is any such structure, regardless of base pair identity, sufficient for enzyme activity? Our and others’ data suggest that while the length and stability of the stem modulate the pseudouridylation rate, the combination of a uridine at the base of a stem-loop structure may be sufficient for PUS1 to recognize it as a target. Second, if guide RNA-independent PUS enzymes recognize and bind to defined RNA structural motifs, and if such motifs are indeed responsible for the majority of mRNA pseudouridylation, are these motifs specifically placed in mRNA to have regulatory functions? Ψs are not randomly distributed within mRNAs, and have been shown to be enriched in the 3’-UTRs and coding region (14), suggesting a functional role. The structural basis for target recognition by PUS1 presented here is a first step towards understanding how PUS enzymes select their targets in mRNA, and if and how this process may be regulated.

## ACKNOWLEDGEMENTS

The authors thank Alexander Zhelkovsky for his pioneering work on PUS1 in the Yigit lab, and Meredith Purchal for helpful comments and discussions. The Berkeley Center for Structural Biology is supported in part by the National Institutes of Health, National Institute of General Medical Sciences, and the Howard Hughes Medical Institute. The Advanced Light Source is supported by the Director, Office of Science, Office of Basic Energy Sciences, of the U.S. Department of Energy under Contract No. DE-AC02-05CH11231. The Pilatus detector was funded under NIH grant S10OD021832. The ALS-ENABLE beamlines are supported in part by the National Institutes of Health, National Institute of General Medical Sciences, grant P30 GM124169

## DATA AVAILABILITY

The structures described in this manuscript have been deposited in the RCSB protein database and are available for public download and examination (PDB ID codes 7R9F and 7R9G). The original source data and raw images corresponding to the biochemical analyses of PUS1 behavior and activity have been uploaded to a public repository (at https://dataverse.harvard.edu/dataverse/scPUS1/) and are also available upon request from the authors.

## Supplemental Materials and Methods

### Electrophoretic mobility shift assays

100 nM (10 pmol) to 1 µM (100 pmol) of the wildtype and the mutant PUS1 enzymes were incubated with 100 nM (10 pmol) of the synthetic RNA oligo R263 in a 100 µl reaction under PUS1 activity assay conditions (see Materials and Methods) for 90 min at 30°C. Immediately before loading onto the gel, 18 µl of each reaction were transferred to a fresh tube containing 2 µl native PAGE loading dye (0.05% xylene cyanol, 50% glycerol). 18 µl of each sample were then loaded on a 6% Novex^TM^ TBE gel that was running at 150 V at 4°C. After 30 min, the gel was stopped and visualized using the Cy2 channel of an Amersham Typhoon Laser Scanner after staining with SYBR Gold (1/20,000 dilution, Thermo Fisher Scientific # S11494) for 10 min.

### Determination of Ψ sites by oligonucleotide UHPLC-MS/MS

3.9 pmol of PUS1-modified and unmodified R397, R398, and R444 oligonucleotides (sequences are listed in **Table S1**) were digested with 1 µL of a 1:50 dilution of RNaseT1 (Thermo Fisher Scientific) in a 30 µL reaction in 1 x NEB buffer r1.1 (10 mM Bis-Tris-Propane-HCl, 10 mM MgCl_2_, 100 µg/ml Recombinant Albumin, pH 7.0) at 37 °C for 30 minutes. RNaseT1 cleavage products were spin filtered at 13,400 rpm for 5 minutes utilizing Ultrafree MC-GV 0.22 µm spin filters (Millipore).

Filtered oligonucleotides were analyzed by UHPLC-MS/MS. UHPLC analysis was performed on a Vanquish Horizon UHPLC (Thermo Fisher Scientific) utilizing an ACQUITY Premier Oligonucleotide C18 Column (Waters) (2.1 x 100 mm, 1.7 µm) with a 23-minute 7% - 35% gradient of solvent A (1% hexafluoroisopropanol (HFIP), 0.1% N,N-diisopropylethylamine (DIEA), 1 μM EDTA) and solvent B (90% Methanol, 10% water, 0.075% HFIP, 0.0375% DIEA, 1 μM EDTA) at a flow rate of 400 µL/min (at 60°C). MS/MS analysis was performed on an Eclipse Fusion Orbitrap Mass Spectrometer (Thermo Fisher Scientific) in data-dependent acquisition mode at a resolution of 60,000 using high energy collision dissociation (HCD) with a stepped normalized collision energy of 22, 24 and 26%.

MS/MS-based oligonucleotide sequencing data were searched utilizing the Nucleic Acid Search Engine (NASE) tool in OpenMS (1) with a 5% false discovery rate and RNase T1 digests of corresponding R397 or R398 oligonucleotide sequences. Pseudo-uridylated oligonucleotides were detected by a procedure adapted from Yamauchi *et al*. (2). Briefly, NASE annotated MS/MS spectra containing a 207.04 m/z doubly dehydrated Ψ nucleoside signature ion and a 211.00 m/z ribose phosphate ion with a mass accuracy tolerance of +/- 0.005 m/z were identified and summed.

### Supplemental Figure Captions

**Fig. S1.**
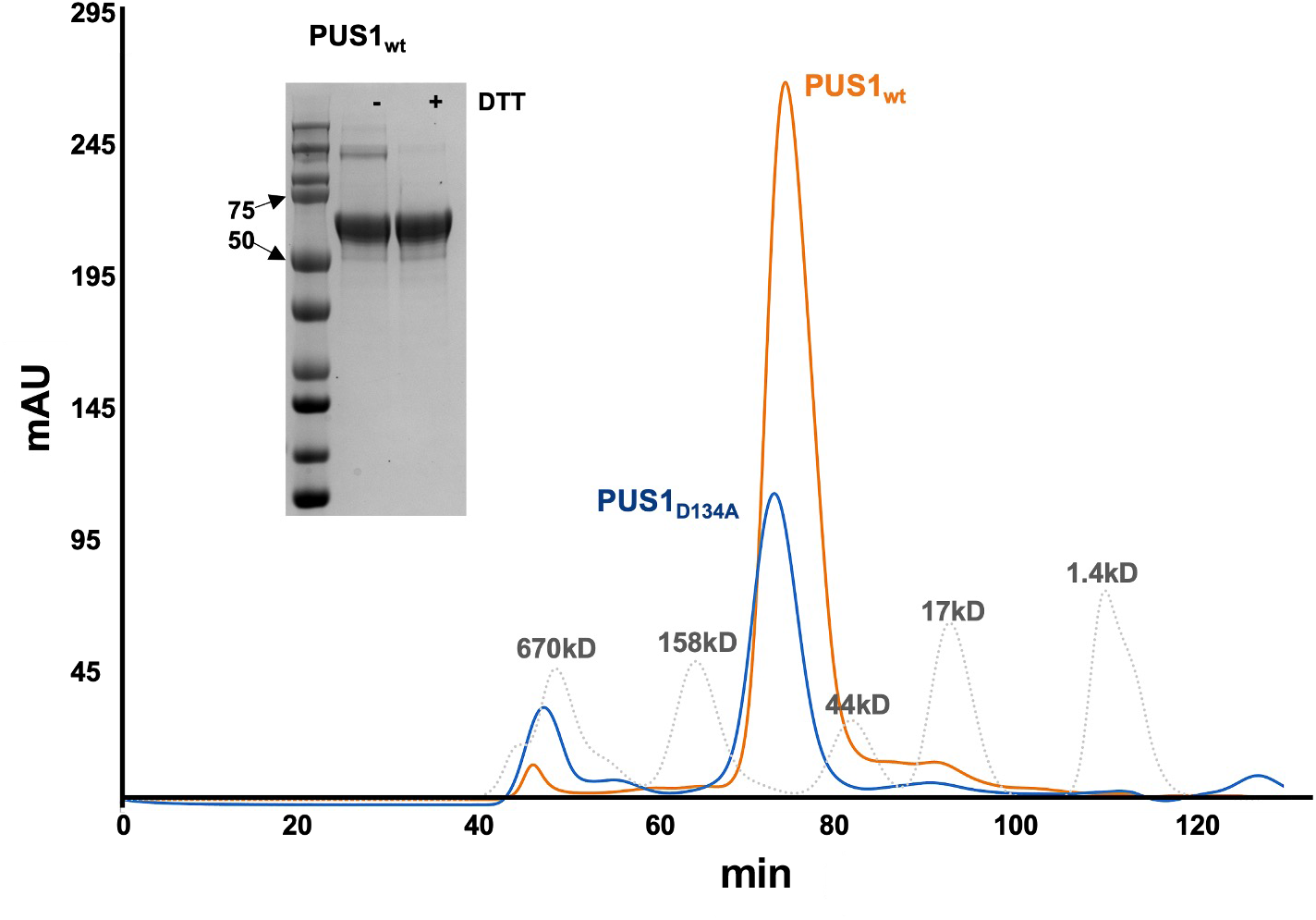
Purification of *S. cerevisiae* PUS1. ***Panel a:*** Wildtype PUS1 (PUS1_wt_, orange) and catalytically inactive PUS1 (PUS1_D134A_, blue) run as monomers on a HiLoad 16/60 Superdex 200 column. Standards (Bio-Rad Gel Filtration Standards) are overlaid in dotted gray and appropriate peaks labeled. ***Inset:*** SDS-PAGE of the final purification product of PUS1_wt_ without (-) or with (+) dithiothreitol.

**Fig. S2.**
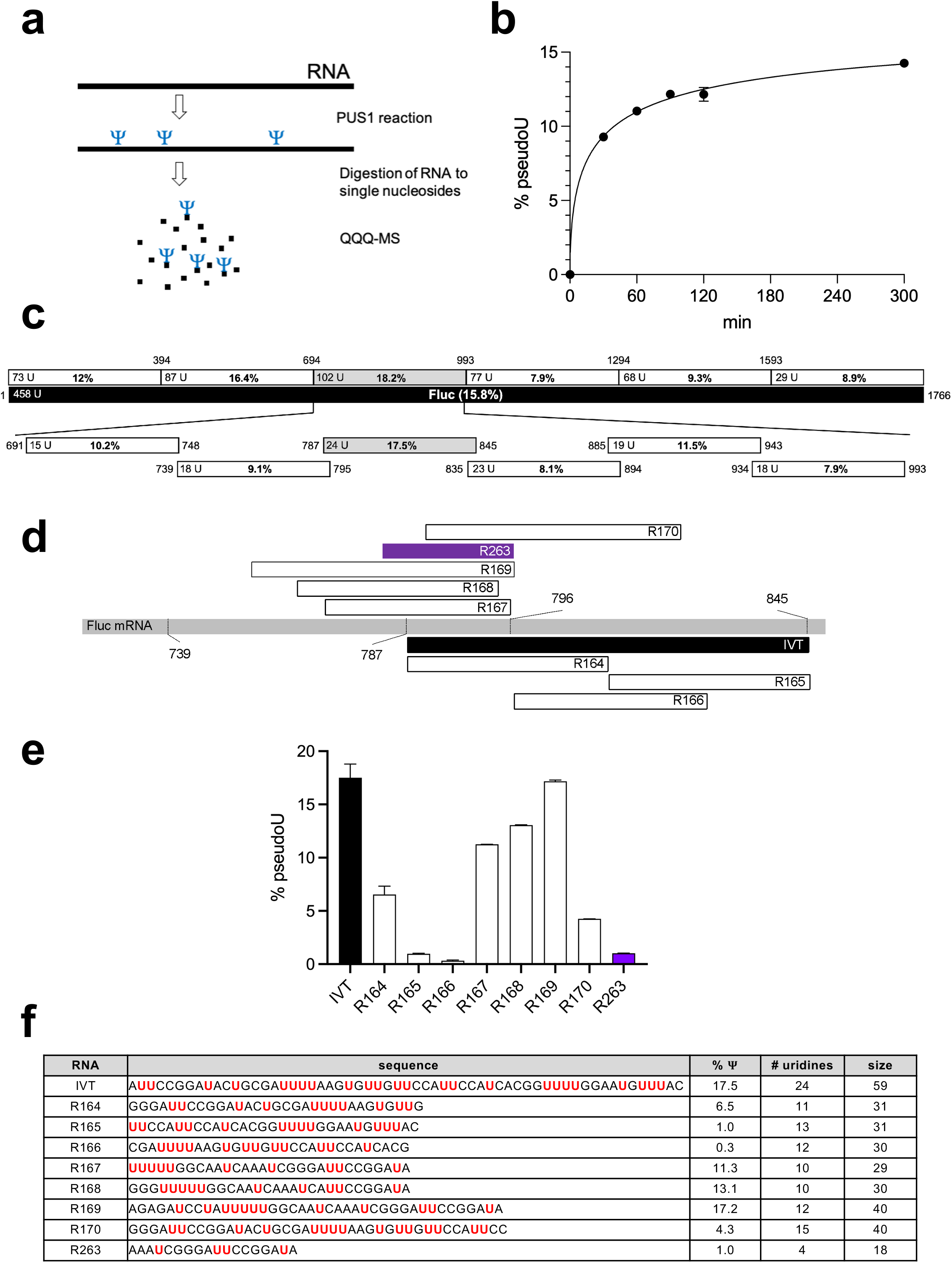
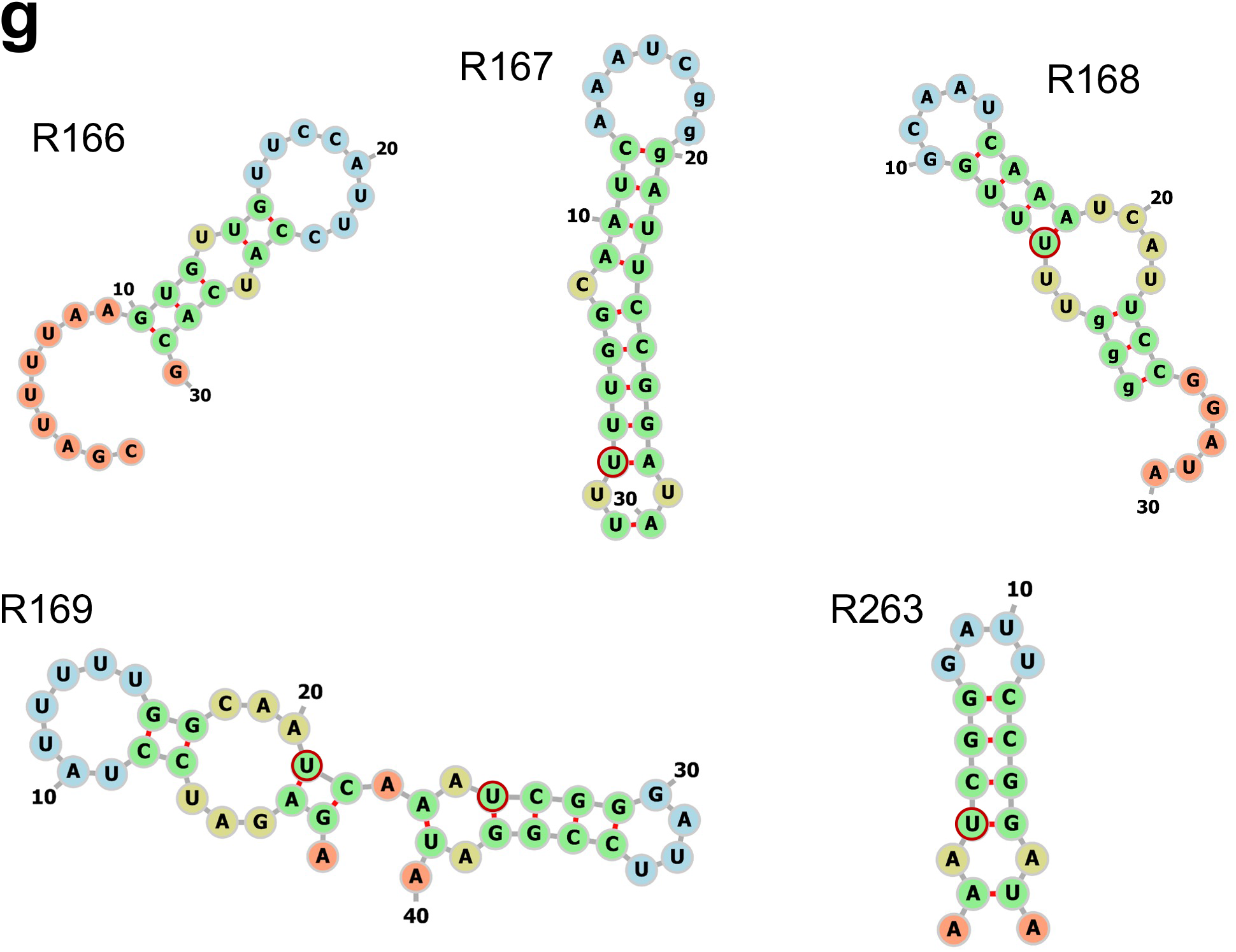
Determination of a suitable RNA substrate for crystallization with PUS1. ***Panel a:*** Schematic representation of the PUS1 activity assay and analysis. ***Panel b:*** PUS1 was incubated with Firefly luciferase (Fluc) mRNA. The graph shows percent of uridine-to-Ψ conversion (% pseudoU) over time in minutes from two independent experimental replicates. Reactions were stopped after 30/60/90/120/300 minutes and analyzed via LC-MS/MS. ***Panel c:*** Schematic view of the RNA substrates generated and tested to determine the optimal RNA substrate for crystallography. The black bar represents the full length Fluc mRNA sequence. The six bars above Fluc represent ∼ 300-mers of the Fluc sequence. The respective start- and end-positions are shown above the segments, the number of total uridines per segment (x U) and the percent of uridine-to-Ψ conversion in a PUS1 reaction after two hours (data from two independent replicas) are shown inside the bars. The bars below Fluc represent ∼ 60-mer substrates between Fluc positions 691 and 993. ***Panel d:*** Schematic view of synthetic RNA oligos and a short *in vitro* transcribed RNA covering Fluc positions 760 to 845. The names of the substrates are shown inside the bars. ***Panel e:*** Percent of uridine-to-Ψ conversion as determined by LC-MS/MS of the RNA oligos by PUS1 after incubation for two hours. Each data set contains at least two replicas. ***Panel f:*** Table showing the sequence of the RNA oligos tested (uridines are highlighted in red), the average percent of uridine-to-Ψ conversion from at least two independent experimental replicates, the number of uridines per RNA oligo, and its respective size in nucleotides. ***Panel g:*** RNA secondary structure predictions of the RNA oligo substrates by MXfold2 (3). Uridines at the base of a stem-loop structure and thus potential PUS1 targets, are highlighted by a red circle.

**Fig. S3.**
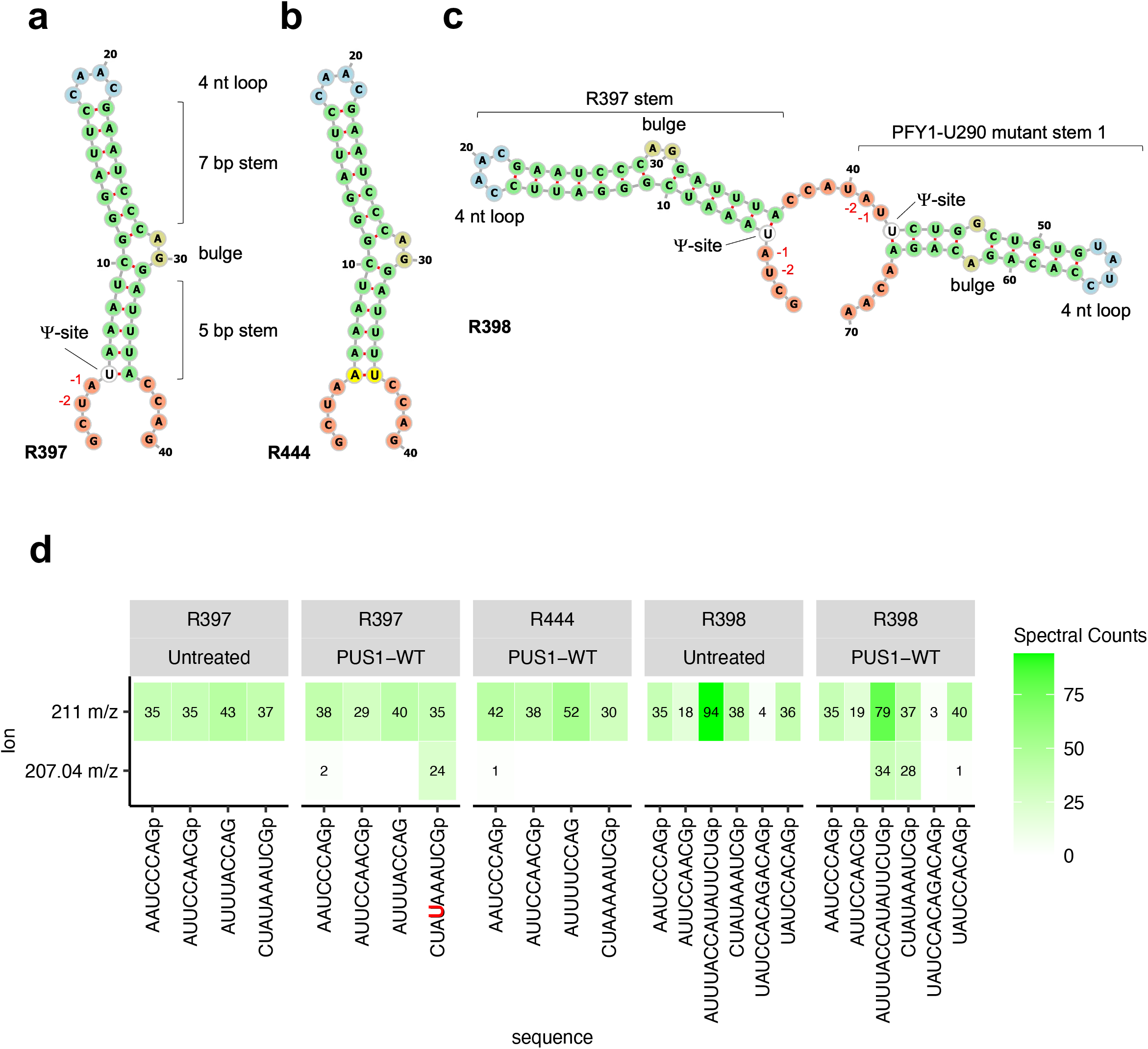
PUS1 pseudouridylates HRU-motif containing RNA stem loop substrates *in vitro*. RNA secondary structure predictions (3) of RNA substrates R397 (***panel a***), R444 (***panel b***), and R398 (***panel c***). The predicted pseudouridylation site, the positions of the H (- 2) and R (- 1) nucleotide in relation to the pseudouridylation site, and RNA structure features are indicated. ***Panel d:*** Heatmaps of the number of oligonucleotide spectra identified by LC-MS/MS analysis in a RNase T1 digest of RNA substrates after incubation with PUS1 indicated above the heatmaps. Shown are the characteristic 207.04 m/z 4′ nucleoside signature ion (bottom) or the 211.00 m/z ribose phosphate ion (top).

**Fig. S4.**
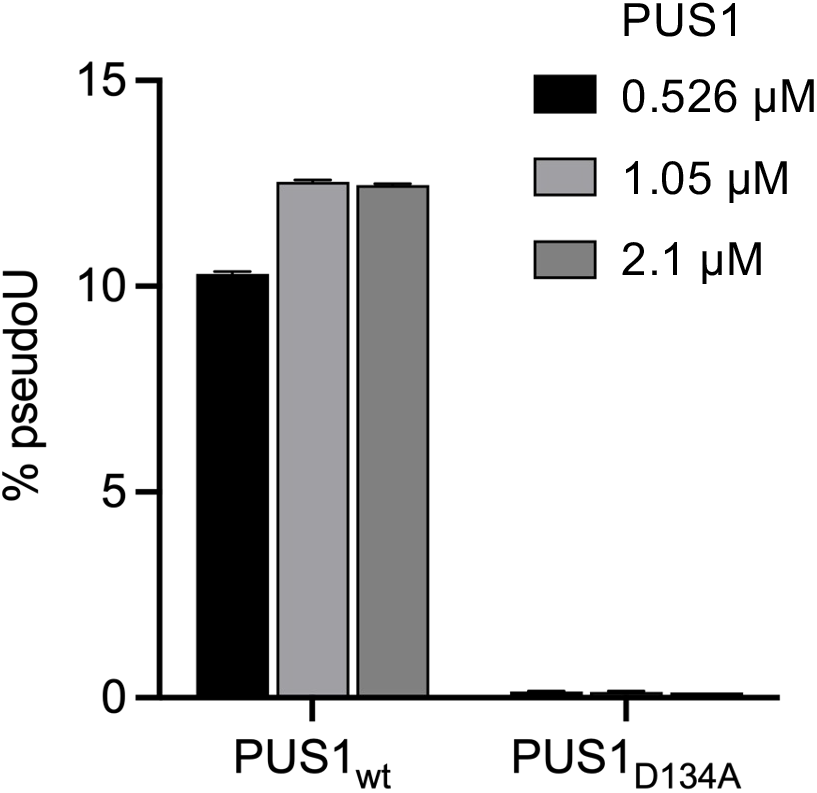
A PUS1_D134A_ enzyme is catalytically inactive. Percent of uridine-to-Ψ conversion as determined by LC-MS/MS of increasing concentrations of PUS1 wildtype (left) and D134A mutant (right) with the RNA oligo substrate R168 after two hours. Data of two replicas are shown.

**Fig. S5.**
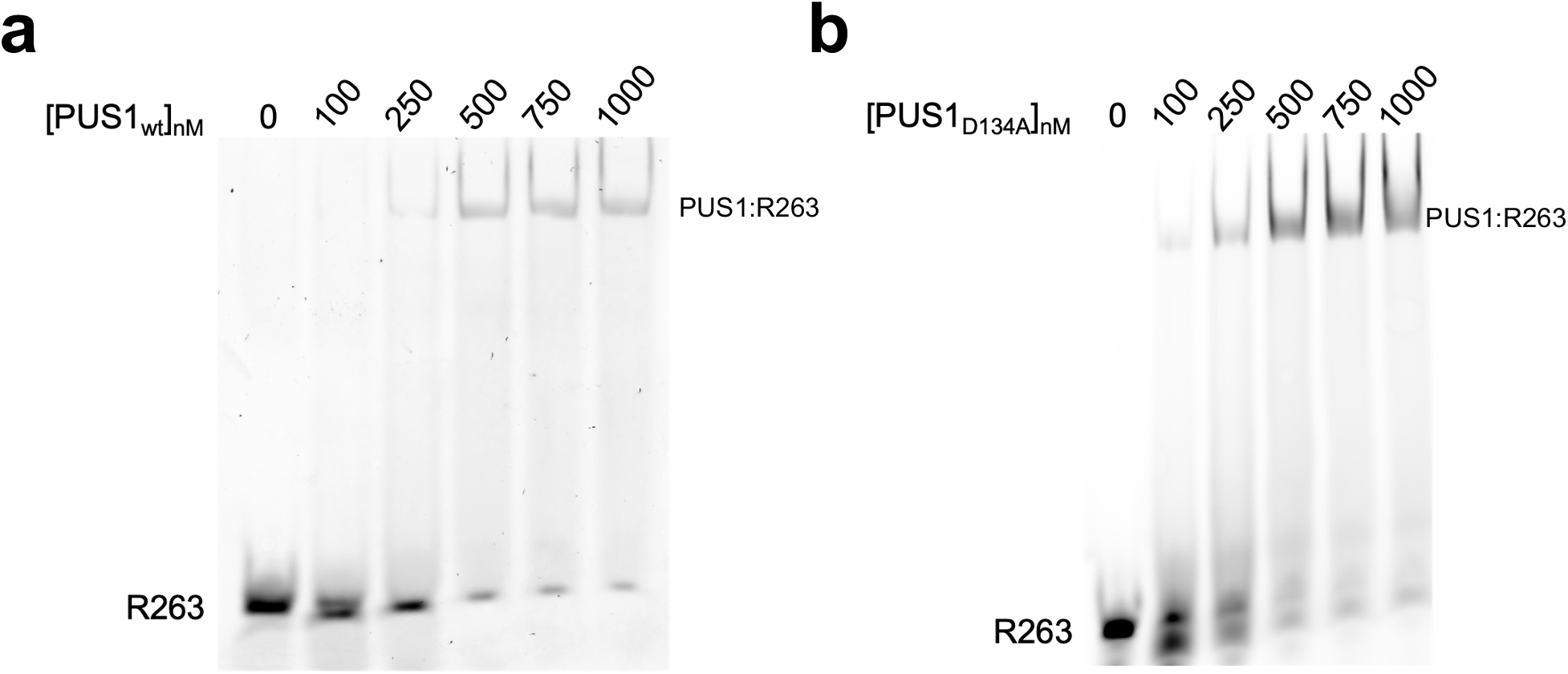
PUS1_wt_ and PUS1_D134A_ stably bind an RNA oligo. Electrophoretic mobility shift assays of increasing concentrations of wild type PUS1 (left) and PUS1_D134A_ (right) as shown above the gels with 100 nM of RNA oligo R263. Free R263 RNA oligo and the PUS1:RNA complex are indicated.

**Fig. S6.**
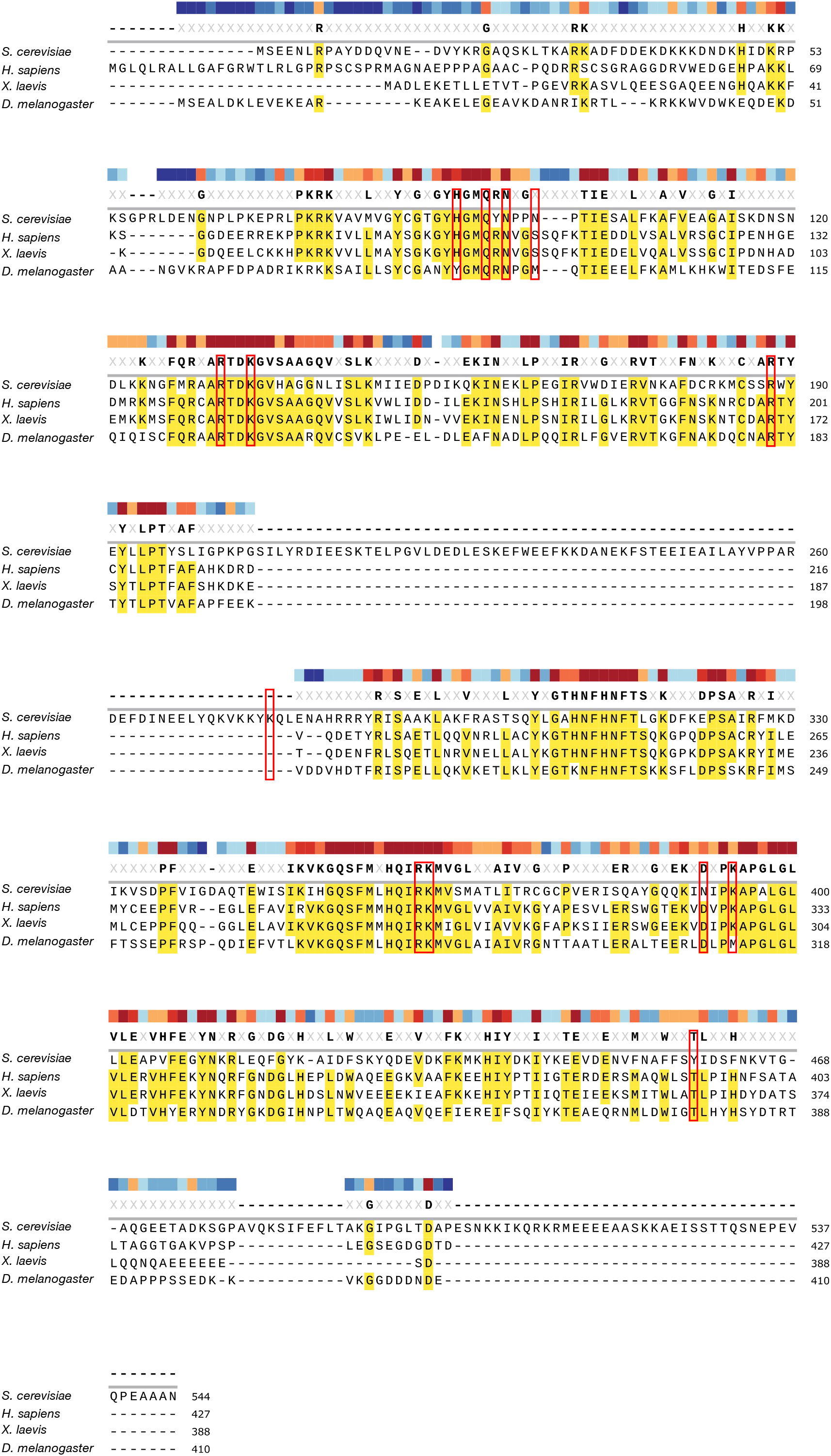
Sequence alignment of eukaryotic PUS1 homologues. Amino acid residues that interact with the substrate RNA are highlighted by red boxes.

**Fig. S7.**
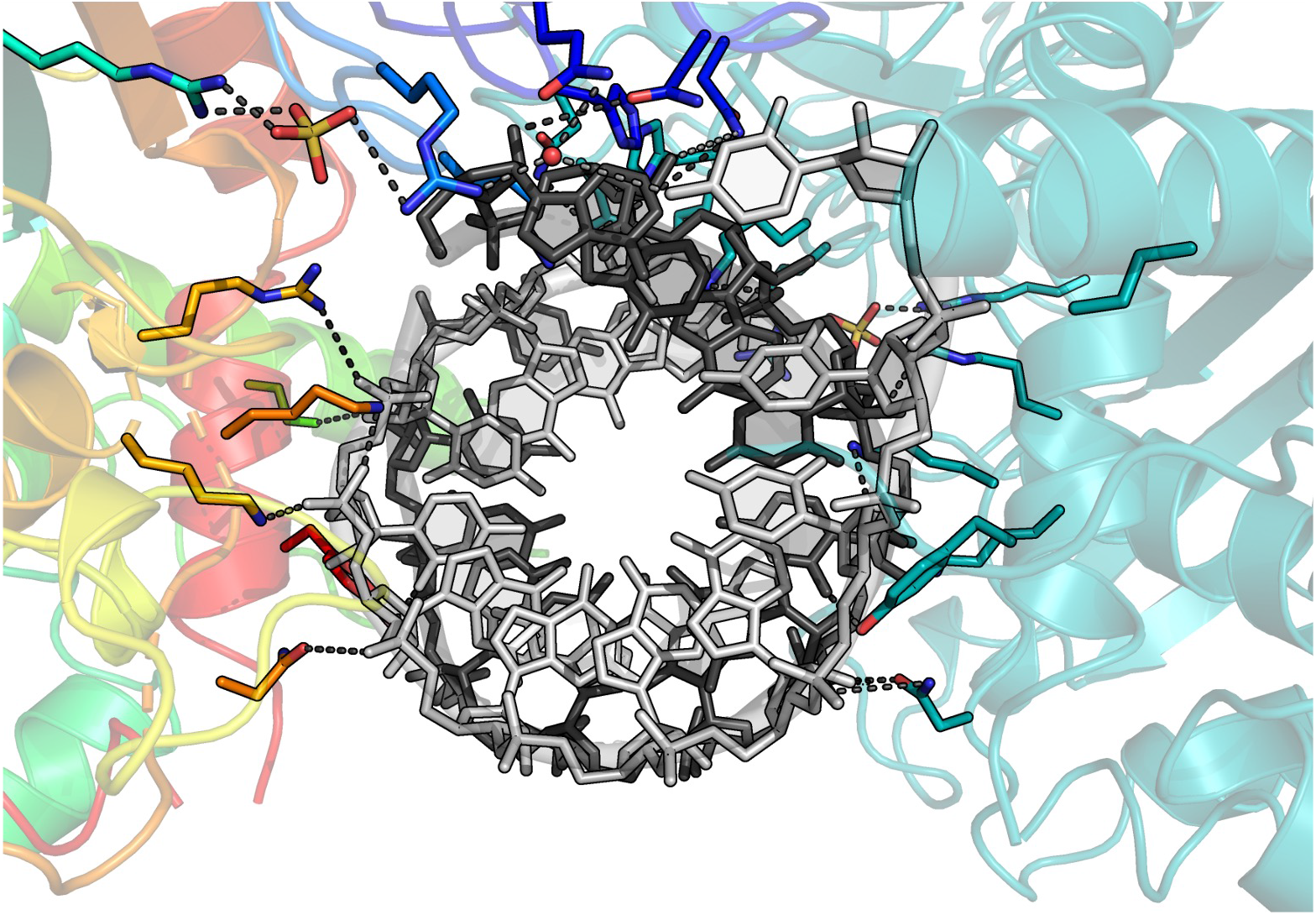
PUS1-RNA contacts. Distribution of contacts around the RNA duplex for both enzyme subunits in the dimeric assemblage.

**Fig. S8.**
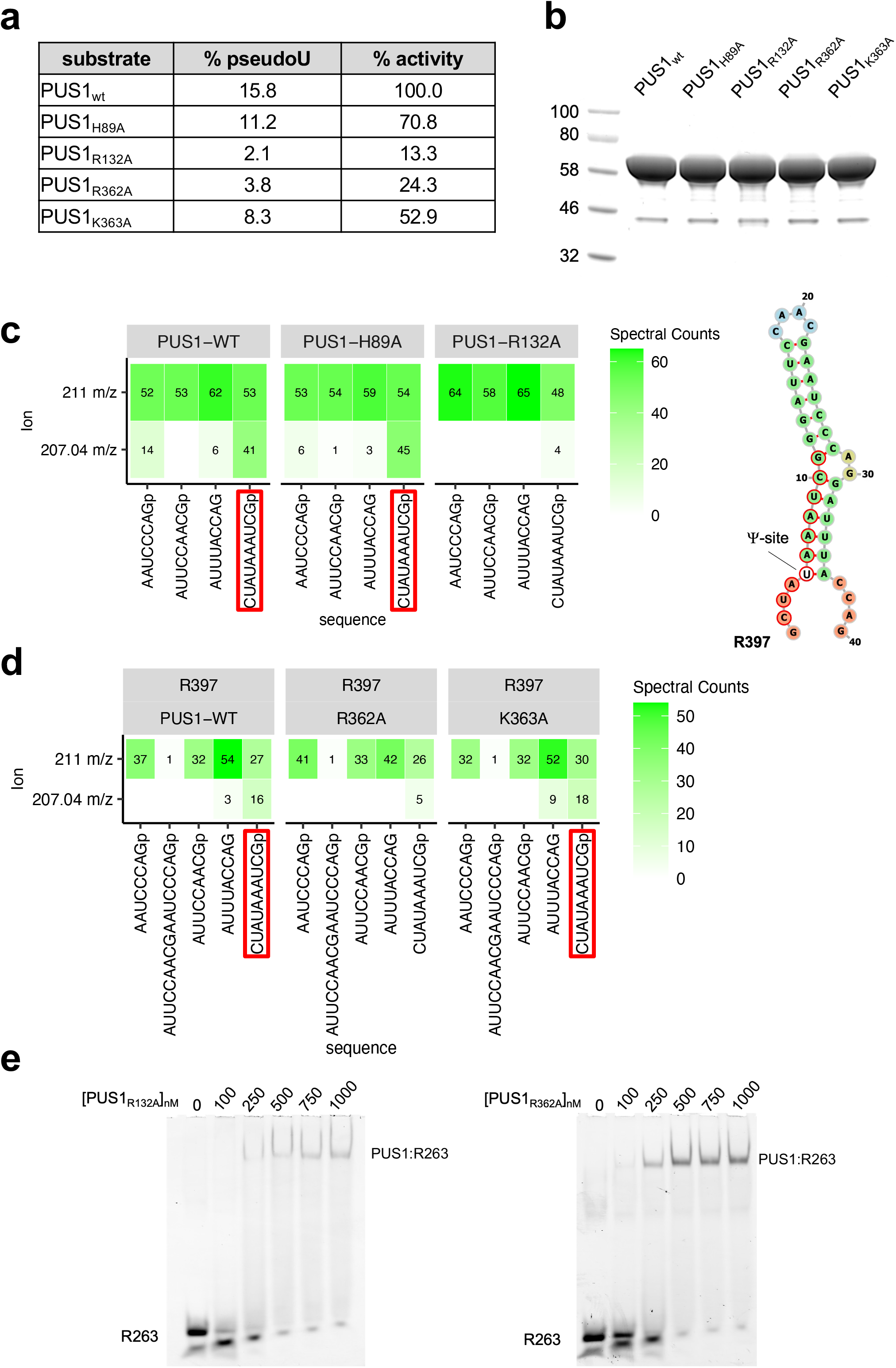
Mutation of RNA-contacting amino acid residues adversely affect PUS1 activity. ***Panel a:*** Table showing the average percent of uridine-to-Ψ conversion of the reactions shown in Figure 3c (from two independent replicates), and the percentage of PUS1 activity, normalized by the percent pseudouridylation in the PUS1_wt_ reaction. ***Panel b:*** Protein gel comparing the input amount and purity of the PUS1 variants in the *in vitro* reactions. ***Panel c:*** Heatmaps of the number of oligonucleotide spectra identified by LC-MS/MS analysis in a RNase T1 digest of RNA substrate R397 after incubation with wildtype PUS1 and cluster A mutants showing either a characteristic 207.04 m/z Ψ nucleoside signature ion (bottom) or a 211.00 m/z ribose phosphate ion (top). The fragments containing Ψ are highlighted with red boxes below the heatmaps and red circles in the RNA structure (right). ***Panel d:*** LC-MS/MS analysis of RNA substrate R397 after incubation with wildtype PUS1 and cluster B mutants, showing the characteristic 207.04 m/z Ψ nucleoside signature ion and the 211.00 m/z ribose phosphate ion. Ψ containing fragments are highlighted with red boxes. ***Panel e:*** Electrophoretic mobility shift assays of the mutant PUS1 enzymes (PUS1_R132A_ left, PUS1_R362A_ right).

**Fig. S9.**
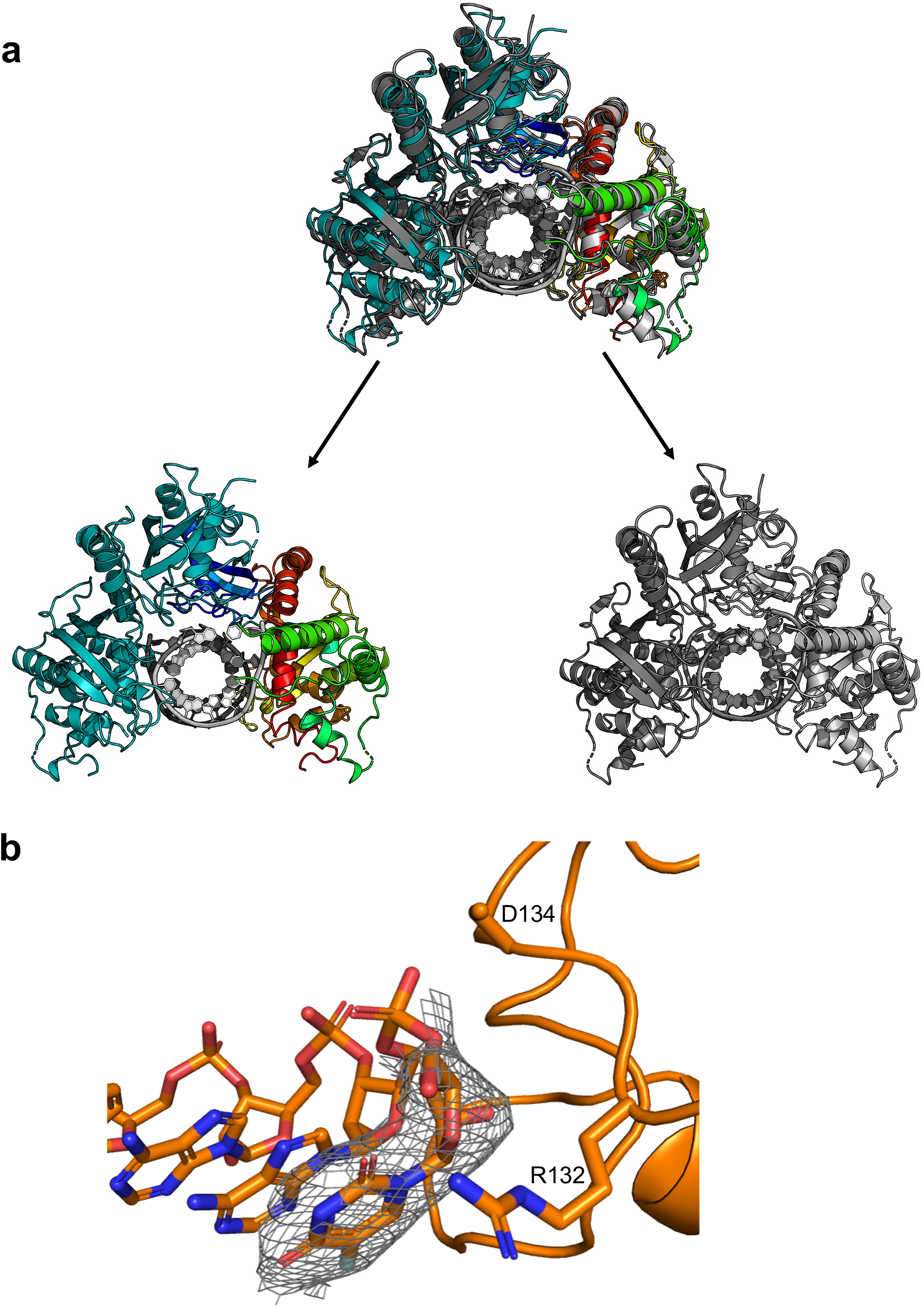
Comparison of PUS1 / RNA complex structures. ***Panel a:*** Superposition and side-by-side comparison of PUS1_D134A_ / RNA complex (colored protein backbones) solved in crystallographic space group C2 and wild type PUS1 / RNA complex (grey protein backbones) solved in an unrelated, P6_1_22 space group. In both structures, two monomers of the enzyme are independently bound in a symmetric arrangement to an RNA duplex that is generated via a crystallographic 2-fold dyad symmetry axis. ***Panel b:*** Model and electron density for wild type PUS1 and the region of the RNA containing the 5’ 5-fluorouracil base. A 2Fo-Fc map (gray) is contoured at 1s. In the corresponding refinement, the base is estimated to display greater than 90% occupancy of the unflipped conformation, as modeled and shown.

**Fig. S10.**
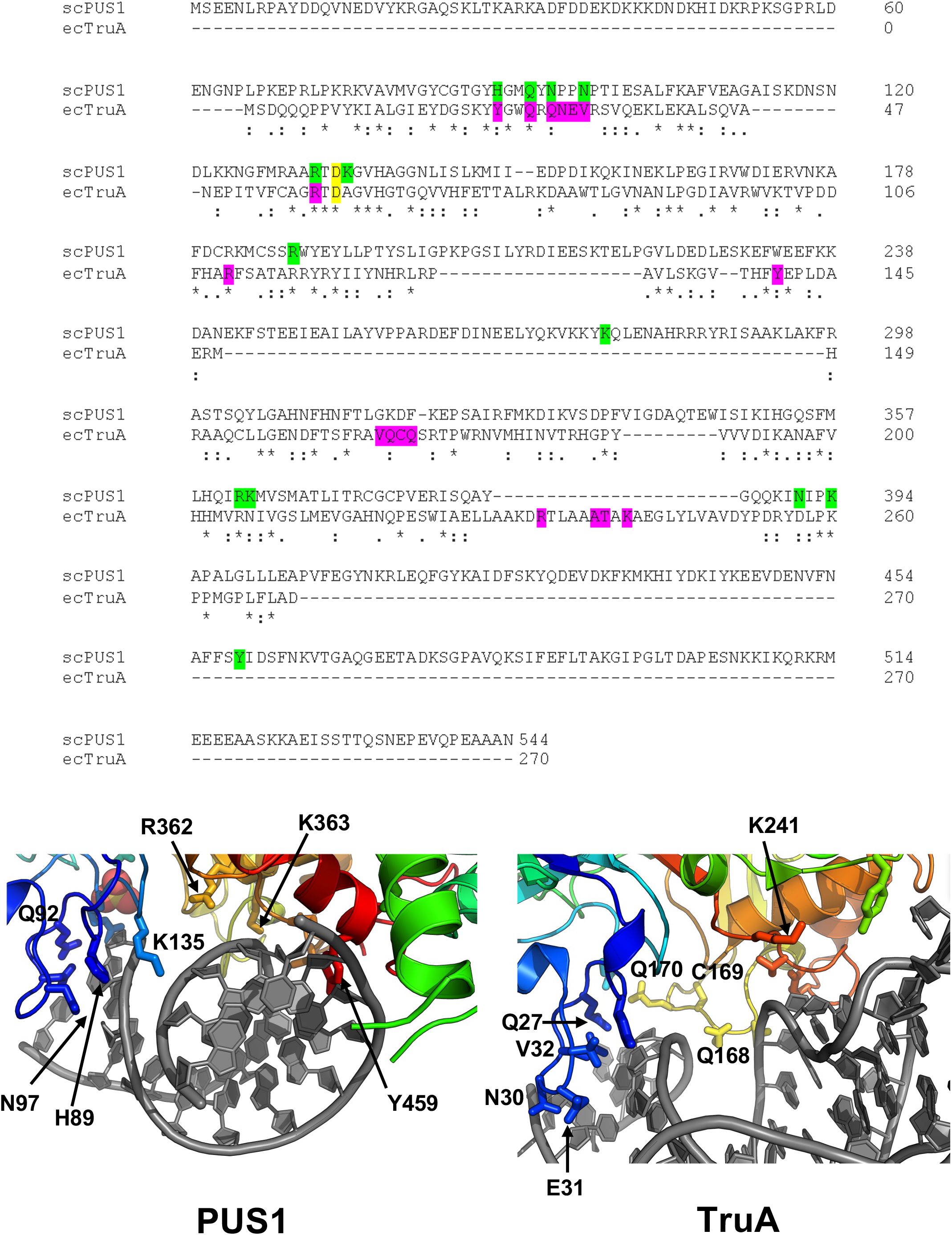
Sequence alignment and RNA interactions of *S. cerevisiae* PUS1 and *E. coli* TruA. The conserved catalytic aspartate at position 134 in PUS1 and position 60 in TruA is indicated by a yellow star and box. RNA-interacting residues are highlighted in green (PUS1) and magenta (TruA) (top panel). Distribution of RNA-contacting residues in the protein-subunit interface for PUS1 (left) and TruA (right; bottom panel).

**Table S1.**
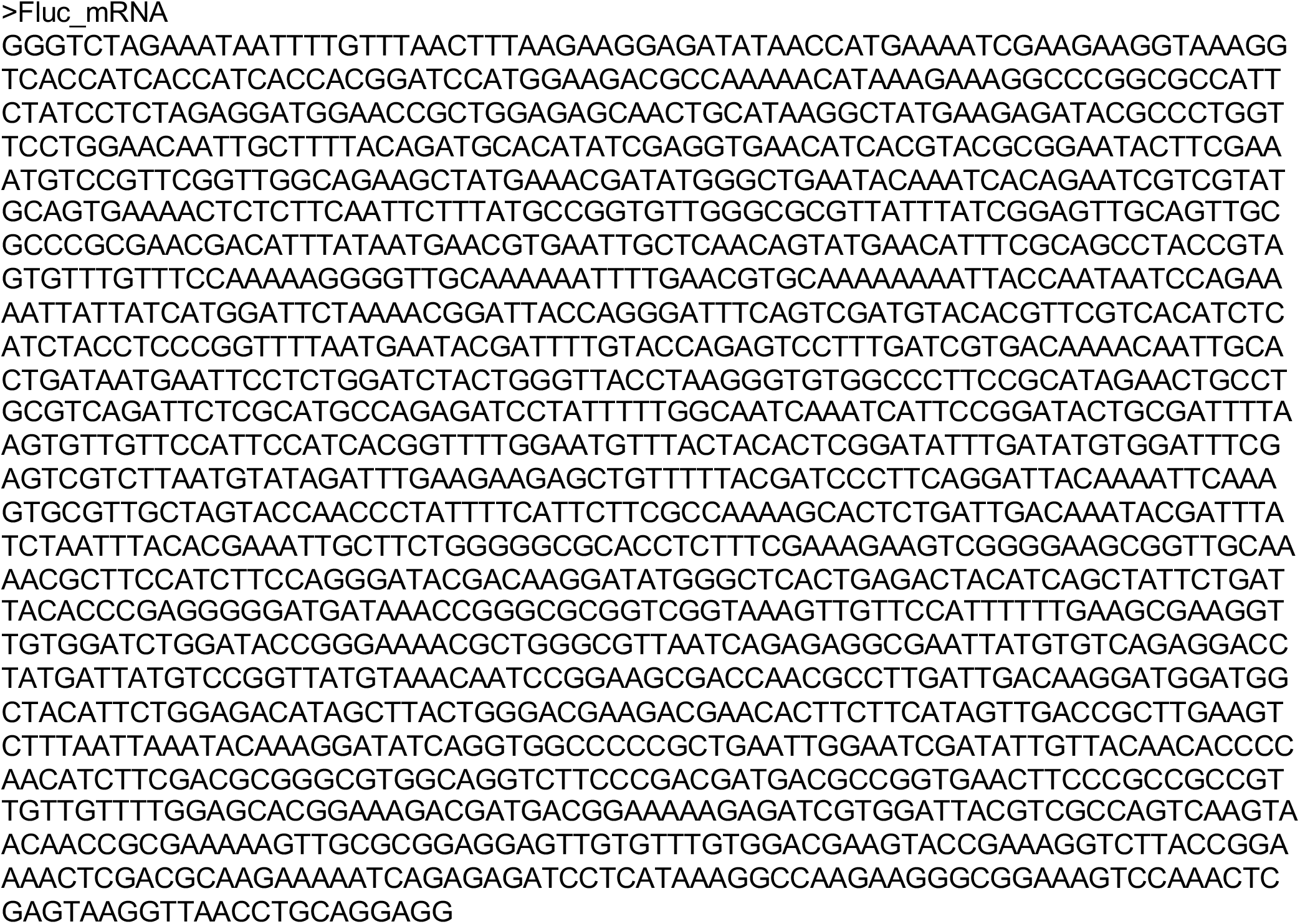
Fluc mRNA sequence.

**Table S2.**
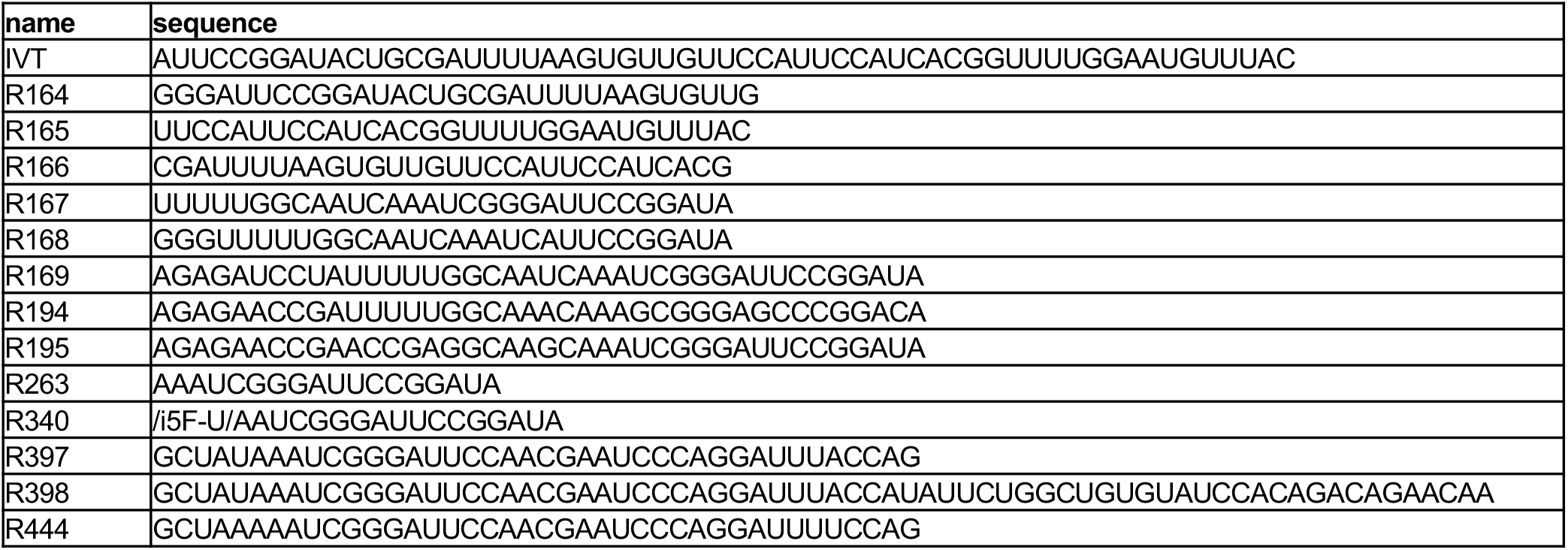
Sequences of the RNA oligos used in this study.

